# Hour-long live-cell TIRF imaging of GFP-talin at ensemble and single-molecule level with low-duty-cycle picosecond supercontinuum excitation

**DOI:** 10.64898/2026.11.23.768601

**Authors:** Felix Margadant, Oddmund Bakke, Michael P. Sheetz, Xian Hu

**Author notes:** Deceased 30 January 2025 (Michael P. Sheetz).

## Abstract

Photobleaching and phototoxicity limit how long living cells can be followed by total internal reflection fluorescence (TIRF) microscopy. We describe an objective-type TIRF microscope illuminated by a picosecond supercontinuum laser spectrally windowed to 20-nm bands, pulse-picked to 1 MHz and delivered through two TIRF illuminators, one per polarisation direction. Single GFP-talin molecules in focal adhesions of living cells were detected throughout uninterrupted recordings of 10–31 min, and drift-corrected, time-stamped localisations resolved adhesion motion of tens of nanometres per minute. In diffraction-limited time-lapse at one frame per second, adhesion brightness, adhesion area, cell footprint and adhesion motility were unchanged after an hour. Under continuous-wave excitation on a comparable commercial TIRF microscope, the same schedule preserved cell behaviour at a low laser setting but adhesions dimmed, and higher settings bleached and retracted cells within a minute of exposure. We provide the datasets and open ImageJ tools for drift correction and localisation.

## Introduction

Photobleaching imposes a practical limit on how often a living cell can be observed by fluorescence microscopy. In camera-based TIRF, where the whole field is illuminated for the full exposure, it determines how many frames a time-lapse can meaningfully consist of, how finely that series can be sampled in time, and whether a single-molecule experiment is possible at all. The limiting factors are the loss of signal through photobleaching and the phototoxicity caused by the reactive products of the same photochemistry. Single-molecule tracking is affected immediately by both and therefore serves as a good test case; similar constraints apply to TIRF imaging, where we test the same illumination method.

Single-molecule localisation microscopy derives its resolution from many sequential images, each containing a sparse set of isolated emitters whose positions are fitted to a few tens of nanometres. In fixed cells the number of frames is limited only by time and by the reservoir of fluorophores; in living cells phototoxicity and the depletion of that reservoir become limiting, while laser power and frame rate need to be high to provide temporal resolution.

Photobleaching has no single mechanism. Irreversible chemistry can start from the excited singlet state, from the triplet state or from radical states, by reaction with molecular oxygen or by absorption of a further photon into higher excited states, and the weight of each route depends on the fluorophore, its environment and the irradiance^1,2^. Long-lived dark states are not in themselves destructive: ground-state-depletion and dSTORM imaging rely on shelving fluorophores reversibly in dark states for milliseconds to seconds^3,4^. What several mitigation strategies have in common is that they keep a molecule that is in an excited or dark state from absorbing again. In scanning microscopy, Donnert et al.^5^ obtained a 5–25-fold increase in the total fluorescence collected before bleaching, for the dye Atto 532 and for GFP, when strong continuous-wave or high-repetition-rate illumination was replaced by pulses separated by more than 1 µs, and attributed the gain to the relaxation between pulses of dark states with lifetimes of about a microsecond, such as the triplet state. For Atto 532, whose triplet lifetime is about 1 µs^5,6^, the explanation is direct. For GFP it cannot be the whole explanation: the triplet state of EGFP is formed with a quantum yield of about 1 % and lives for about 5 ms at room temperature (4.2 ms by phosphorescence decay), three orders of magnitude longer than such a pulse interval, and it is this state that leads into oxidative photochemistry^7^. Boudreau et al.^8^ found on a confocal microscope that 250-ps pulses at 20–80 MHz halved the bleaching rate of EGFP relative to continuous illumination at equal exposure, independently of the repetition rate, and that microsecond pixel dwell times repeated by line averaging reduced it nine-fold. The second effect is compatible with a millisecond triplet state, because the interval between successive visits of the beam is of that order, as in the proposal that fast scanning reduces photobleaching^9^ and in the reduction obtained with resonant scanning in STED microscopy^6^; the first, like the GFP result of Donnert et al., is not explained by it. Photobleaching of fluorescent proteins is in addition supra-linear in irradiance^10^, and depopulating the triplet state with near-infrared light reduces it^11^, so both the rate at which a molecule is excited and the time it spends in the triplet state matter. Reviews of phototoxicity list pulsed illumination among the few generally applicable mitigations^12^. Whether such a benefit carries over to camera-based TIRF microscopy, where the whole field is illuminated for the full camera exposure, has to our knowledge not been reported. A picosecond supercontinuum laser is a convenient test bed for this question. Its pulses are far shorter than the excited-state lifetime of GFP, and at the pulse energies used here a molecule is excited by a given pulse with a probability of only a few per cent (Methods). A pulse picker sets the interval between pulses from 50 ns (20 MHz) to 1 µs (1 MHz) and beyond; even the shortest of these intervals is many times the excited-state lifetime, so the interval matters only for longer-lived states such as the triplet state. The 1 µs interval used here is the pulse separation above which Donnert et al. observed their gain for GFP.

Focal adhesions couple the actin cytoskeleton to the extracellular matrix through a layered set of adaptor proteins, of which talin is the primary integrin-to-actin linker. Fluorescent speckle microscopy has shown that actin flows retrogradely over adhesions and that adhesion proteins move with it to different degrees: F-actin moves at ∼25 nm s⁻¹ in the distal lamellipodium but slows to ∼2 nm s⁻¹ in the lamella^13^; within focal adhesions, talin and vinculin flow at 0.1–0.3 µm min⁻¹, whereas paxillin and integrins move at less than 0.1 µm min⁻¹^14,15^. This hierarchy of speeds is the signature of the molecular clutch, and single-molecule speckle microscopy indicates that only 4.1 % of talin molecules are linked to actin and to the substrate at the same time^16^. Talin in adhesions is therefore a demanding test specimen for a long-duration single-molecule method: the molecules are immobilised for variable times, exchange with a large cytosolic pool, and the biologically interesting motions are tens of nanometres per minute.

Here we combine a supercontinuum excitation laser with a standard objective-type TIRF microscope (Figure 1) and record GFP-tagged talin in living cells at one frame every two seconds for 10–31 minutes (Figure 2). We characterise the persistence of single-molecule detection and the precision with which slow adhesion motion can be measured, and we provide the analysis as ImageJ plugins. We then show, in an independent dataset recorded on the same instrument, that the same illumination supports diffraction-limited time-lapse of GFP-talin for a full hour at one frame per second with no loss of adhesion brightness, adhesion area, cell footprint or adhesion motility (Figure 3). Finally, we compare this with continuous-wave excitation at the same acquisition schedule (Figure 4): at a low laser setting the cells remained healthy for an hour but their adhesions dimmed, and at higher settings they bleached and retracted within a minute of cumulative exposure. The continuous-wave recordings were made on a commercial TIRF microscope built for that purpose and not on the pulsed instrument, where continuous-wave laser lines coupled into the two-polarisation illumination path gave poor, uneven images. The comparison therefore gives continuous-wave excitation its favourable case, and we state where the difference in specimen and the absence of a power calibration limit the conclusions.

**Figure 1.**
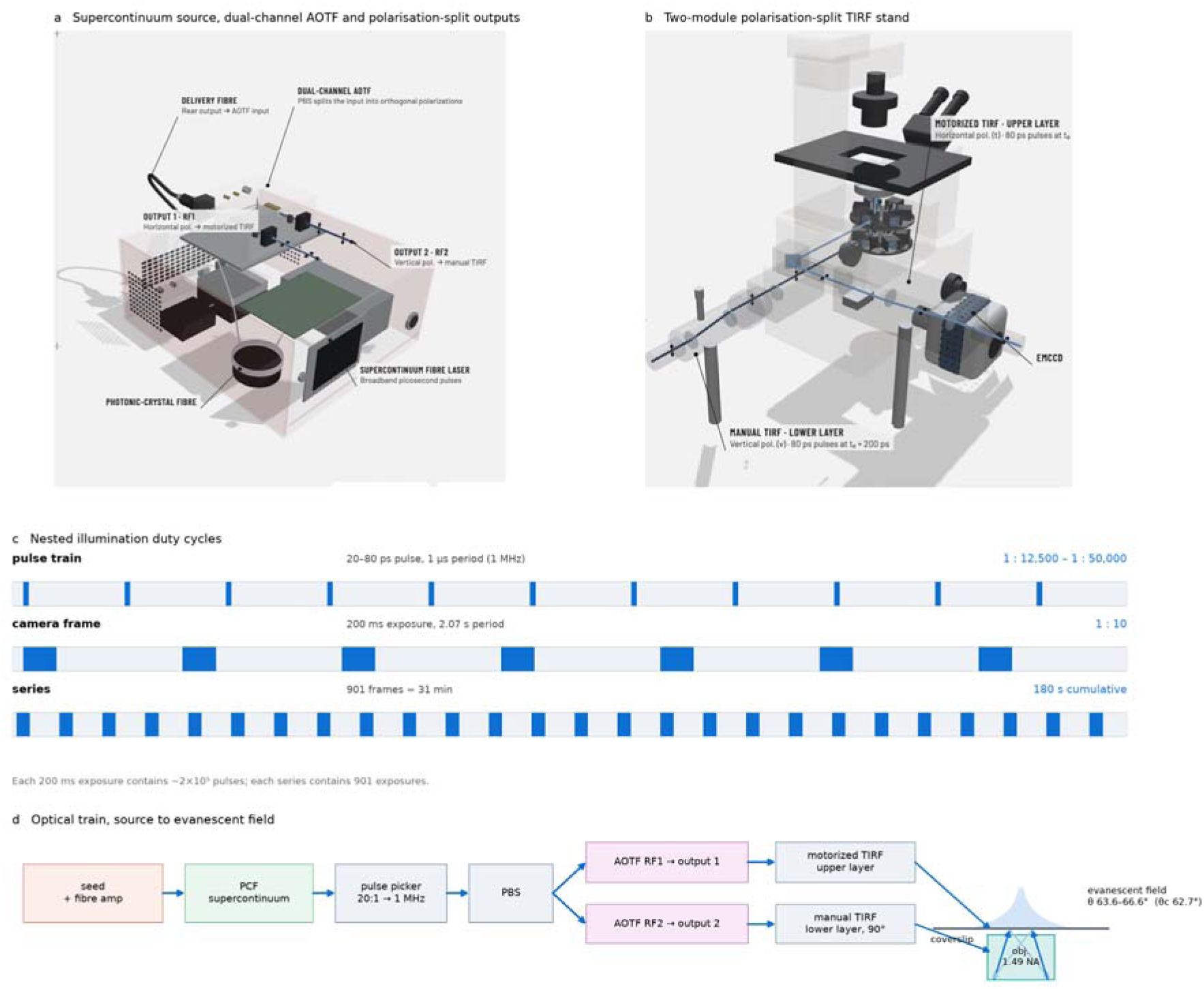
Low-duty-cycle pulsed TIRF illumination: source, beam path and timing. (a) Supercontinuum fibre laser and dual-channel acousto-optic tunable filter. Broadband picosecond pulses generated in the photonic-crystal fibre leave by the rear delivery fibre and enter the AOTF enclosure, where a polarising beamsplitter separates the input into two orthogonal linear polarisations; each is diffracted by its own acousto-optic crystal (radio-frequency drives RF1 and RF2) and leaves by its own output, output 1 (horizontal) feeding the motorised TIRF illuminator and output 2 (vertical) the manual one. Enclosure walls are drawn transparent. (b) The two-module polarisation-split TIRF stand. The two illuminators occupy the two layers of the Ti-E filter turret — motorised on the upper layer, manual on the lower — and deliver their beams to the objective back focal plane at azimuths 90° apart; emission is collected by the same objective onto the EMCCD. (c) The nested illumination duty cycles. Top, the pulse train: a 20–80 ps pulse every 1 µs after 20:1 pulse picking. Middle, the camera frame: a 200 ms exposure every 2.07 s, 1:10. Bottom, the series: 901 such exposures, 31 min of elapsed time containing 180 s of cumulative excitation. Each 200 ms exposure contains ≈ 2 × 10⁵ pulses. Bands are schematic and not to scale within a row. (d) The optical train from the seed oscillator to the evanescent field, with the angle of incidence used (63.6–66.6°; critical angle 62.7°) and the exponential intensity profile of the evanescent field above the coverslip. Panels a, b and d are renderings of the instrument as built in 2017; photographs of the same components are in Figure S3.

**Figure 2.**
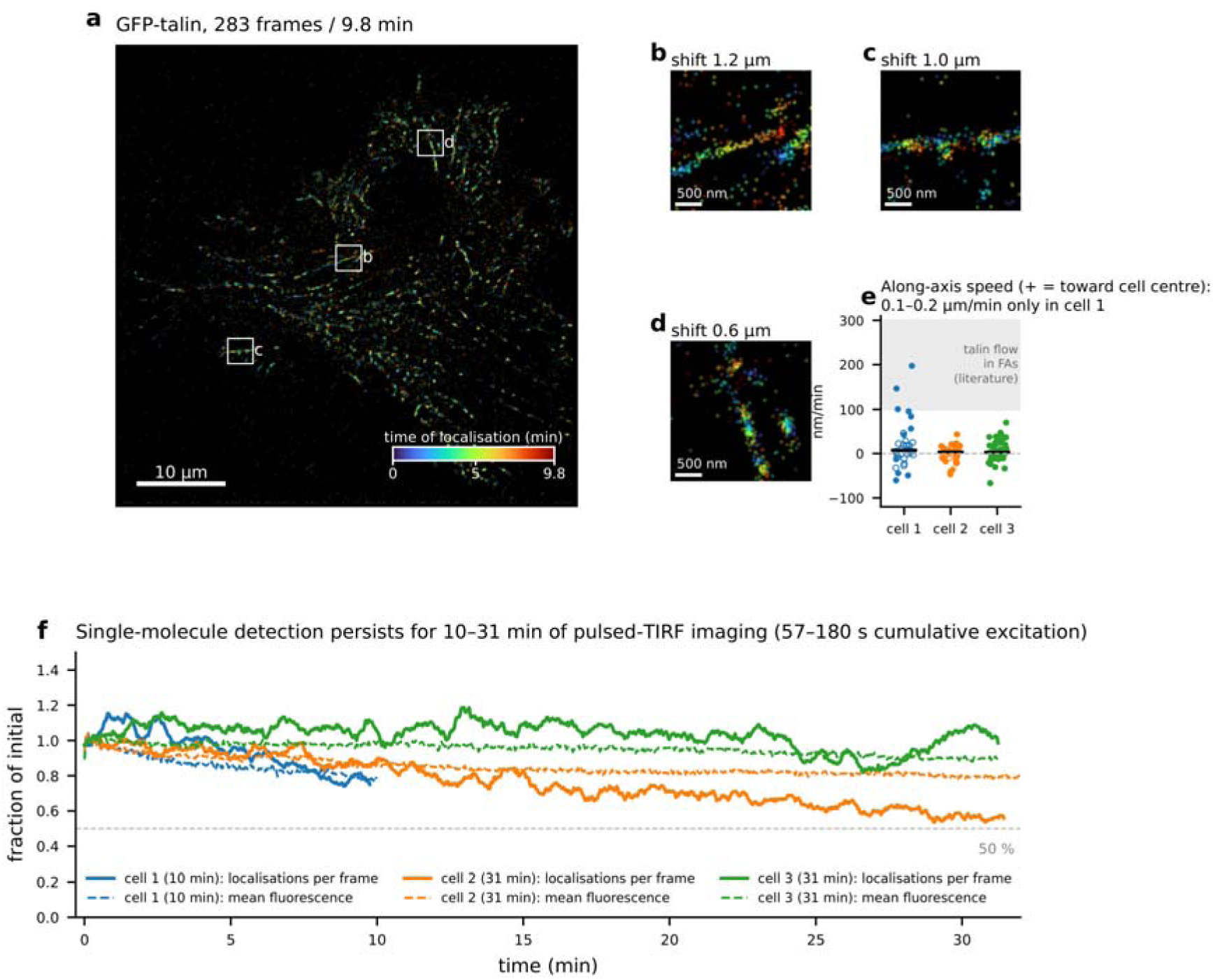
Pulsed-TIRF live-cell SMLM of GFP-talin resolves 10–31 minutes of adhesion dynamics. (a) 32,271 localisations from 283 frames (9.8 min) rendered at 20 nm per pixel and coloured by the time of localisation (colour bar). Scale bar 10 µm. (b–d) 3 µm regions around the three adhesions with the largest early-to-late centroid displacement, rendered at 10 nm per pixel; the ordered colour gradient along each adhesion reports its translocation. Scale bars 500 nm. (e) Along-axis speed of individual adhesions (localisations projected on the adhesion axis oriented toward the cell centre, Theil–Sen slope versus time; positive = retrograde) in cells 1–3 (n = 36, 35, 52 adhesions with ≥ 150 localisations over ≥ 6 min); filled symbols, slope significant at p < 0.01; bars, medians; grey band, 0.1–0.3 µm min⁻¹ range reported for talin flow in focal adhesions. (f) Localisations per frame (solid, 15-frame running mean) and mean background-subtracted GFP channel fluorescence of the cell (dashed) for the three cells, each normalised to the first frames; dotted line, 50 %.

**Figure 3.**
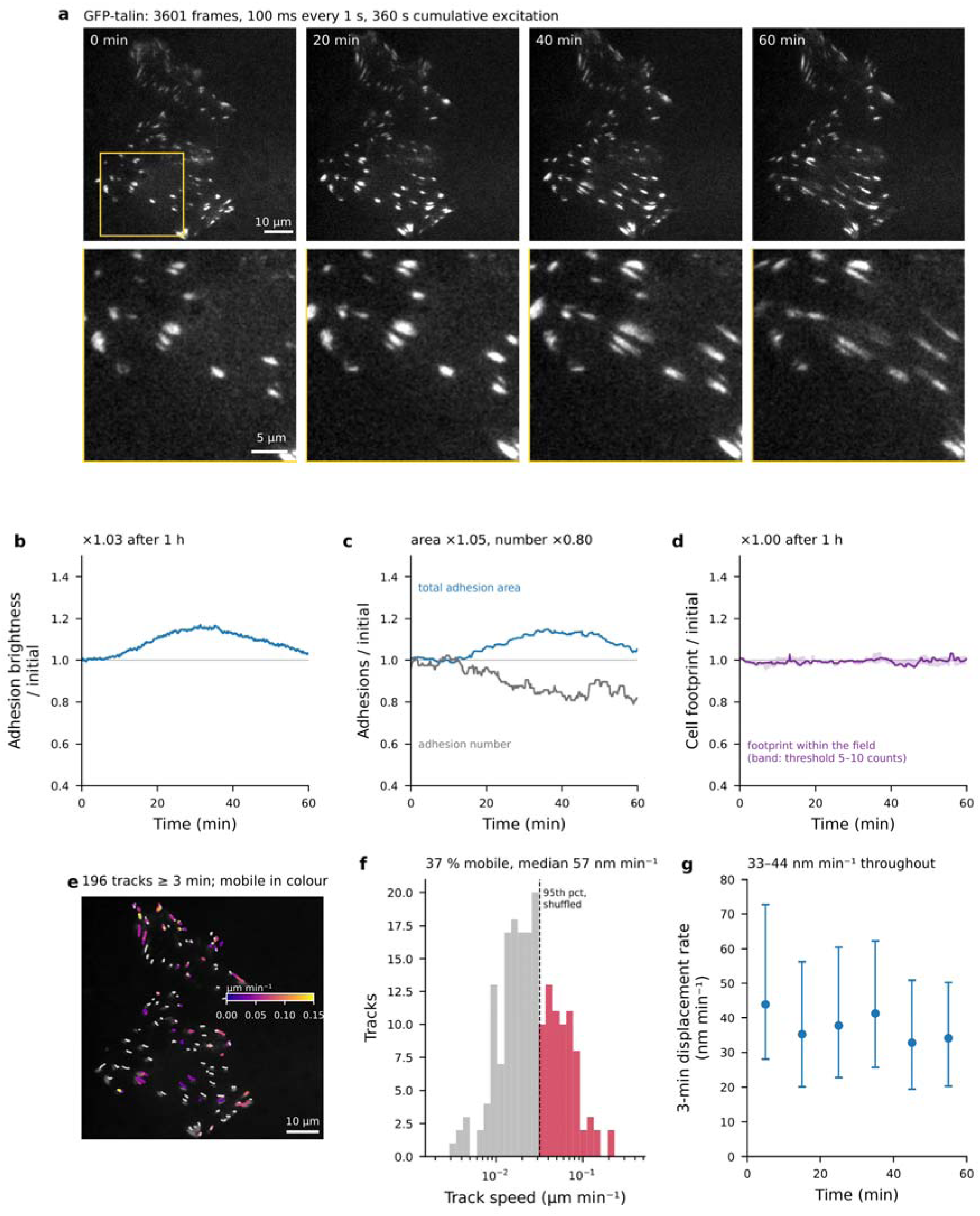
One hour of pulsed TIRF at one frame per second leaves GFP-talin brightness, adhesion area, cell footprint and adhesion movement unchanged. MDA-MB-231 cell co-expressing GFP-talin and paxillin-mApple (GFP channel only acquired), 3601 frames, 100 ms every 1 s, 360 s of cumulative excitation, AOTF line 1 at 50 %, EM gain 300, 160 nm per pixel. (a) GFP-talin after 0, 20, 40 and 60 min; bottom row, the boxed region enlarged. Each image is the mean of five consecutive frames, smoothed with a Gaussian of σ = 0.7 pixel, at the contrast of the first image; no drift correction (drift < 1 pixel over the hour). Scale bars 10 µm (top) and 5 µm (bottom). The full hour is shown in Video S2. (b) Adhesion brightness (99th percentile of the frame minus external background), normalised to the first 1 % of frames; running median of 15 frames. (c) Total area (blue) and number (grey) of segmented adhesions, on 5-frame means every 10 s, normalised to the first 2 min. (d) Cell footprint within the field (diffuse GFP-talin signal > 7 counts above the per-frame background after smoothing; band, thresholds of 5 and 10 counts), normalised to the first 2 min. The cell extends beyond the right and lower edges of the field. (e) Adhesion centroid tracks of at least 3 min over the first image, dimmed; tracks faster than the shuffled-time threshold of (f) in colour by speed, the others in grey. (f) Track speeds (slope of a linear fit of position against time); dashed line, 95th percentile of the same tracks with time points shuffled (32 nm min⁻¹). (g) Median and interquartile range of 3-min displacement rates of all tracked adhesions in successive 10-min intervals; includes the centroid noise of stationary adhesions. Lines in (b–d) at unity. The other hour-long recordings and the dual-colour control are in Figure S5; values for all acquisitions are in Table S3. Source data: Data S1.

**Figure 4.**
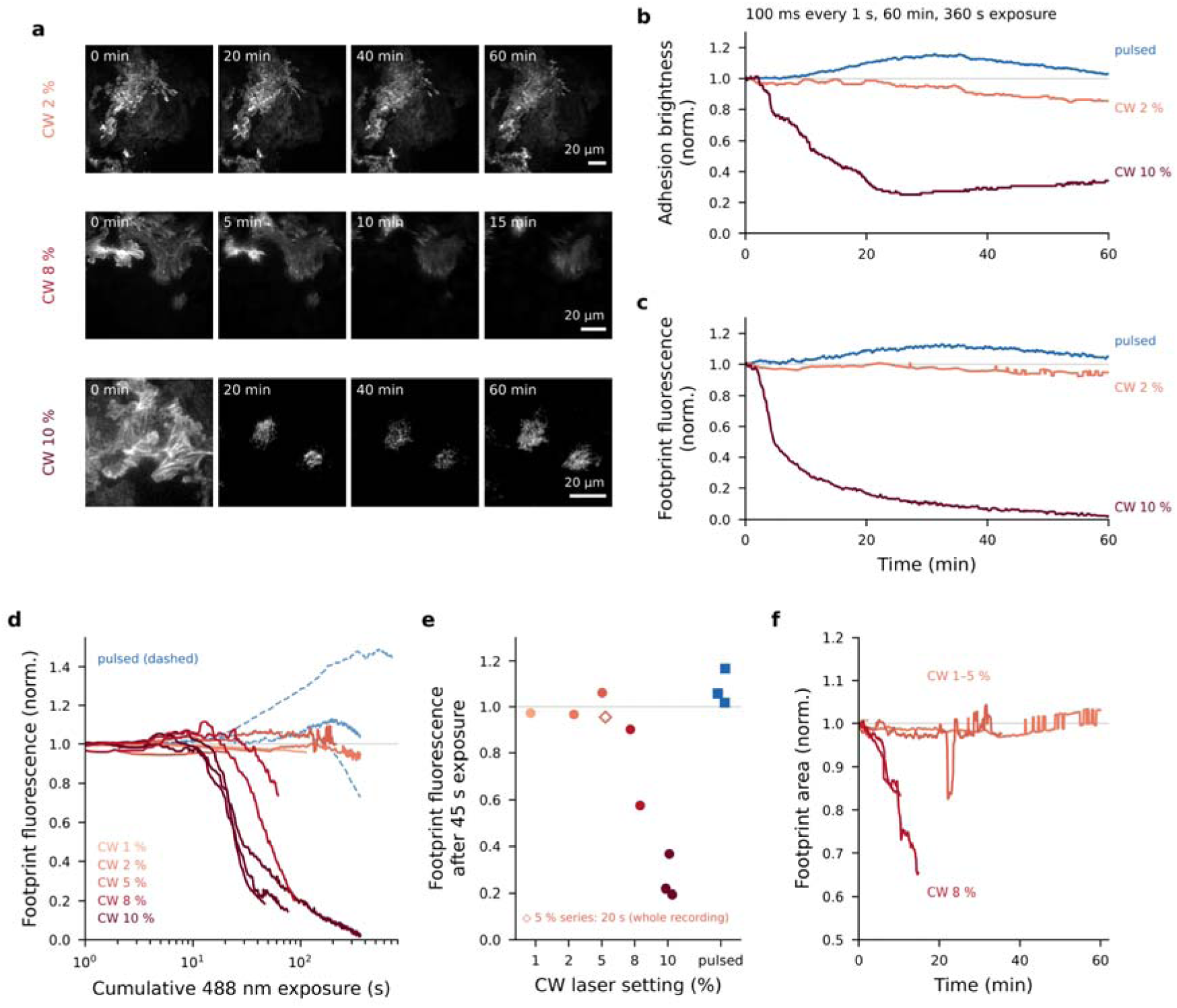
Continuous-wave TIRF power series. U2OS cells expressing vinculin-GFP imaged on a Nikon Ti2-E H-TIRF microscope (Apo TIRF 100×/1.49, Prime 95B sCMOS) with continuous-wave 488 nm excitation at laser settings of 1–10 %, 100 ms exposures every 1 s (1 % series: streamed at 0.117 s per frame); recorded 2026-11-06 to 2026-11-11 (Table S4). (a) Frames of three series at the times shown, each row at the contrast of its first frame: 2 %, 60 min; 8 %, 15 min; 10 %, 60 min (the 2 % and 10 % series at the schedule of the hour-long pulsed series). Images 2× binned and smoothed (Gaussian, σ = 1 pixel). Scale bars 20 µm. (b) Adhesion brightness (99th percentile of the frame minus external background, normalised to the first 1 % of frames; the Figure 3 read-out) versus time for the two 60-min continuous-wave series and the pulsed series of Figure 3 (blue), all 100 ms every 1 s; the 2 % series is analysed in full in Figure S4. (c) Footprint fluorescence of the same series: background-subtracted fluorescence integrated over the initial cell footprint for the continuous-wave series, fixed-mask mean for the pulsed series. (d) Footprint fluorescence versus cumulative 488 nm exposure (logarithmic axis) for all continuous-wave series, coloured by laser setting, and the hour-long pulsed series (dashed; Figure 3 and Figure S5a). (e) Footprint fluorescence after 45 s of cumulative exposure, per recording; open diamond, a 5 % series at 20 s, its whole recording. (f) Footprint area of the cells that could be segmented throughout: 1–5 % series (thresholded per frame relative to the frame’s own brightness) and isolated cells at 8 % (per-frame Otsu threshold; Methods). The 10 % fields (confluent, or with segmentation unstable once bleached) are not plotted. The continuous-wave and pulsed recordings differ in cell line, labelled protein, camera and instrument, and the laser power at the sample was measured for neither; the pulsed series are shown for scale. Running medians of 15–25 frames. Source data: Data S1.

## Results

### A supercontinuum, pulse-picked, polarisation-split TIRF illuminator

The excitation source is a picosecond supercontinuum fibre laser whose broadband output is spectrally windowed by a four-channel acousto-optic tunable filter and pulse-picked from its 20 MHz fundamental down to 1 MHz, giving 20–80 ps pulses separated by 1 µs (Figure 1a, c; Methods). Because the supercontinuum is unpolarised or only poorly polarised, it is split into its two orthogonal linear components rather than passed through a polariser, and the two beams are coupled into two TIRF illuminators mounted on the two layers of the filter turret at azimuths 90° apart, so that the full in-band power reaches the sample (Figure 1b; Figure S3). One design feature is not evident from the layout: the fibres that carry the two polarisations differ in length by 3 cm, so the two pulses reach the sample about 150 ps apart. The pulses, 20–80 ps long, therefore do not overlap in time, yet both arrive well within the excited-state lifetime of virtually all fluorophores. The probability that a molecule in the ground state is excited is therefore independent of the orientation of its transition dipole in the plane of the coverslip and is set by its absorption cross-section, which gives very homogeneous images. From the specification of the source, a GFP molecule is excited by a given pulse with a probability of about 1–3 % (Methods, equations 1–3), far from saturation, so the fluorescence is expected to scale linearly with pulse energy. The complete path from the seed oscillator to the evanescent field is shown in Figure 1d.

### Pulsed TIRF sustains single-molecule detection of GFP-talin for 10–31 minutes

Cells expressing talin tagged with GFP at its N-terminal head (and with mCherry at its C-terminus, recorded in a second channel) were imaged at one frame pair every 2 s for 10–31 min (acquisition parameters in Methods and Table S1). Individual frames contained isolated, diffraction-limited spots on a diffuse background (Figure 2a; Video S1), and detection followed by Gaussian fitting returned 32,000–54,000 localisations per recording, 48–114 per frame. Detection persisted: the number of localisations per frame at the end of the series was 56–104 % of its initial value, and the mean GFP fluorescence of the cell 78–90 % (Figure 2f). Because talin turnover and adhesion remodelling change both quantities as well — one recording showed no decline at all over 180 s of cumulative excitation — these losses are upper bounds on photobleaching. The mCherry channel bleached faster than GFP in the same recordings (Notes S1 and S3), as mCherry generally does; the smaller gain for red fluorescent proteins is discussed under Limitations of the study. Three quarters of the localisations fell inside adhesions, which occupied a third of the cell footprint, and most were seen in a single frame: a detection is a brief bright period of an immobilised molecule (≈ 0.35 s in a series streamed at three frames per second), so localisation counts are detection events and not numbers of molecules (Note S1). Stage drift, up to 1.3 µm in the first series, was measured by phase cross-correlation of ten-frame averages and removed to a residual of 15–29 nm r.m.s., below the single-frame localisation precision (Methods; Figure S1g–i).

### Time-resolved localisation resolves stationary and mobile adhesions

Rendering all localisations colour-coded by acquisition time gives a map of the adhesion field in which each adhesion carries its own 10- or 31-minute history (Figure 2a–d; Figure S1a–c). Most adhesions appear as mixed-colour structures: talin was localised at the same positions throughout the recording. A minority show an ordered colour gradient along their long axis, which means that the cloud of localisations moved during the recording (Figure 2b–d). Projecting the localisations of each adhesion onto its long axis and regressing position against time quantifies this (Methods): in one recording a third of the adhesions moved significantly, most of them toward the cell centre at 0.06–0.2 µm min⁻¹, whereas in the others motion was resolved in more adhesions but was slower, about 20 nm min⁻¹ (Figure 2e; Table S1). These speeds lie within the range reported for talin and vinculin flow in focal adhesions and for lamellar actin retrograde flow, and an order of magnitude below lamellipodial actin flow^13−15^. A plausible reading is that the mobile adhesions are those in which talin is coupled, through a slipping clutch, to the actin network, whereas the stationary majority are anchored to the substrate. The measurement is of the population — it does not distinguish the flow of talin through a stationary adhesion from sliding of the adhesion itself — but it shows that motion of tens of nanometres per minute can be followed in single-molecule data for half an hour in one uninterrupted recording (Note S2).

### Cells tolerate 15–20 min of pulsed TIRF

To test whether such recordings perturb the cells, further cells were imaged in both channels every 5 s for 15–20 min, an interval suited to a diffraction-limited read-out of cell behaviour (Figure S2; Table S2). The cells kept their shape: the footprint changed by −16 to +8 % (median −1.5 %), and none rounded, blebbed or detached. The number of adhesions was constant and their total area changed by a median of −4.5 % (one cell retracted a rim and lost 23 %). Adhesions turned over normally, with new adhesions assembling in every cell. GFP fluorescence was retained at 0.90–1.02 of its initial value (median 0.94), mCherry at 0.75–0.94 (median 0.79) and the number of single-molecule detections per frame at 0.68–1.10 (median 0.88). The 2-s series, re-analysed in the same way, lost more adhesion area (22–47 %); the two sets differ in exposure, gain and duty cycle as well as in cell state, so the cause is not identified, and a phototoxic contribution at the shorter interval cannot be excluded (Note S3).

### GFP-talin survives an hour of pulsed TIRF at one frame per second

The series above are limited to 31 min. To ask how far the photostability extends, we analysed an independent dataset recorded on the same microscope in MDA-MB-231 cells co-expressing GFP-talin and paxillin-mApple: diffraction-limited time-lapse recordings of 4 min to 8 h at frame intervals of 1–10 s (Table S3). We say that a recording was survived if three things hold at its end: the fluorophore has survived (the brightness of the adhesions is retained), the structures have survived (the adhesions are still present and occupy the same area), and the cell has survived (its footprint and the motility of its adhesions are unchanged). A recording at one frame per second for an hour meets all three criteria (Figure 3; Video S2). Adhesion brightness ended at 1.03 of its starting value (Figure 3b). The adhesions remained individually resolved and remodelled throughout: their total area ended at 1.05, while their number fell to 0.80 as adhesions elongated and merged (Figure 3a, c). The cell footprint within the field was unchanged (1.00; Figure 3d). About a third of the adhesions (37 %) moved, most of them toward the cell centre, at a median of 57 nm min⁻¹ (Figure 3e, f) — within the range published for sliding focal adhesions^17,18^ — and the rate of adhesion displacement did not slow over the hour (Figure 3g), as it would if the cell were progressively photodamaged.

The other hour-long recordings bracket this one, ending at 1.37 and 0.54 of their starting adhesion brightness, and the longest recording, 8 h at a 2 % duty cycle, ended at 0.46 (Figure S5a; Table S3). A declining trace cannot by itself be attributed to photobleaching, because focus drift, adhesion disassembly, protein turnover and migration all reduce the signal; an undiminished trace, by contrast, cannot be produced by a bleached fluorophore population. The two-channel recordings provide an internal control: paxillin-mApple ended below its starting brightness in every recording, whereas GFP-talin ended brighter in several, and in some it rose while mApple fell in the same frames, which rules out changes of focus or of the evanescent field as the cause of the mApple loss (Figure S5b, c; Note S4).

### The same schedule under continuous-wave excitation: cells stay healthy at a low laser setting, but adhesions dim

No continuous-wave time series was recorded on the pulsed instrument, to which we no longer have direct access. To place the pulsed recordings against conventional illumination we therefore used a commercial TIRF microscope (Nikon Ti2-E with H-TIRF, Apo TIRF 100×/1.49 objective, Prime 95B sCMOS camera; objective numerical aperture and camera quantum efficiency comparable to those of the pulsed instrument) and U2OS cells expressing vinculin-GFP, and applied the acquisition schedule of the hour-long pulsed recording — 100 ms exposures every 1 s, a frame-level duty cycle of 10 % — at 488 nm laser settings of 1–10 % (Figure 4; Figure S4; Table S4). The read-outs were those of Figure 3, with the analysis parameters scaled to the smaller pixel (Methods); the implementation reproduces the tabulated values of the 2018 series analysed from raw files (0.53, 1.38 and 1.03 against 0.54, 1.37 and 1.03).

At a 2 % setting a cell was followed for the same hour (3600 frames, 360 s of cumulative excitation; Figure S4a). By the criteria of cell behaviour it was unperturbed: its footprint was unchanged (×1.00; 0.99–1.00 over a two-fold range of threshold; Figure S4e), and the rate of adhesion displacement was constant over the hour (62–69 nm min⁻¹ in each 10-min interval; Figure S4f). Its adhesions, however, dimmed. Adhesion brightness held at 0.99 of its initial value for the first 20 min and then declined, to about 0.95 at 30 min, 0.89 at 45 min and 0.85 at 60 min (Figure S4b), and the total area of the adhesions segmented at one fixed threshold fell to 0.76 (Figure S4c). The number of segmented adhesions fell to 0.77 while their mean size did not change (1.35 µm² at the start, 1.33 µm² at the end; Figure S4d), which is the pattern expected if adhesions fade below a fixed threshold rather than being remodelled. In the pulsed recording the number of adhesions fell by a similar factor (0.80), but the adhesions that remained grew (from 1.5 to 1.9 µm²), and brightness and total area ended at 1.03 and 1.05 (Table 1). The two recordings therefore agree in the two read-outs that report cell behaviour and differ in the two that depend on the survival of the fluorophore.

**Table 1.** The schedule-matched hour-long recordings under pulsed and continuous-wave.

|  | Pulsed (Figure 3) | Continuous-wave, 2 % (Figure S4) |
|---|---|---|
| Specimen | MDA-MB-231, GFP-talin | U2OS, vinculin-GFP |
| Instrument and camera | supercontinuum TIRF (2018), EMCCD | Nikon H-TIRF (2026), sCMOS |
| Schedule | 100 ms every 1 s, 3601 frames | 100 ms every 1 s, 3600 frames |
| Cumulative excitation (s) | 360 | 360 |
| Single-frame adhesion signal-to-noise ratio | 10 | 56 |
| Adhesion brightness, end / start | 1.03 | 0.85 |
| Total adhesion area, end / start | 1.05 | 0.76 |
| Adhesion number, end / start | 0.80 | 0.77 |
| Mean adhesion size, start → end (µm²) | 1.46 → 1.91 | 1.35 → 1.33 |
| Cell footprint, end / start (range over thresholds) | 1.00 (0.98–1.02) | 1.00 (0.99–1.00) |
| 3-min displacement rate, range over six 10-min intervals (nm min<sup>−1</sup>) | 33–44 | 62–69 |
| Adhesion tracks ≥ 3 min | 196 | 1971 |

End / start ratios are the means of the last and first 1 % of frames (brightness) or of the last and first 2 min (segmentation read-outs). The displacement rates include centroid noise, which differs between the two recordings with adhesion density, and are compared for their constancy over the hour, not for their absolute value.

The two recordings differ in more than the light source: cell line and labelled protein, camera, instrument and year. The single-frame adhesion signal-to-noise ratio was higher in the continuous-wave recording (56 against 10), so by that measure the continuous-wave image was not the more weakly excited of the two; with different cameras this does not establish the relative irradiance, and the laser power at the sample was measured on neither instrument. The dimming also began only after 20 min, whereas first-order photobleaching under constant illumination begins at once, so a slow biological change in the vinculin-GFP adhesions is not excluded. We conclude that continuous-wave TIRF at a low laser setting sustains an hour of imaging at this schedule with a measurable loss of adhesion signal that the pulsed recording did not show, and that the present data cannot attribute the difference to pulsing.

### Higher continuous-wave settings bleach and retract the cells within a minute of exposure

At 1–5 % the cells kept most of their GFP signal (footprint fluorescence 0.94–1.01 at the end of the recordings) and none retracted. At 8–10 % the outcome was different (Figure 4): footprint fluorescence held for the first 16–57 s of exposure and then fell, reaching half its starting value after 25–50 s in most recordings, and the cells retracted — isolated cells that could be segmented throughout ended at 0.66 and 0.83 of their starting area. Because retraction lifts fluorescence out of the evanescent field, the loss of TIRF signal at these settings combines photobleaching with the loss of cell–substrate contact. The loss rose much more steeply than the laser setting: at 10 % the fluorescence halved after 25–29 s of exposure, whereas at 2 % it stood at 0.94 after 360 s, where proportionality to setting × exposure would predict a halving after 125–145 s. On this microscope, then, the useful duration of continuous-wave GFP imaging was limited by the laser setting rather than by cumulative exposure (Table S4; Note S5).

## Discussion

### What the data establish

Three results stand on the data. First, with picosecond supercontinuum excitation pulse-picked to 1 MHz, single GFP-talin molecules were detected for 10–31 min at a frame-level duty cycle of about 10 %, and for 10 min in a stream of frames without gaps; the number of detections per frame at the end of the recordings was 0.56–1.04 of its initial value in the 2-s series and 0.68–1.10 in the 5-s series. Second, with the same source an ordinary diffraction-limited time-lapse of GFP-talin ran for an hour at one frame per second and ended with adhesion brightness, adhesion area, cell footprint and adhesion motility where they started (Figure 3). Third, at the same schedule under continuous-wave excitation the cell remained healthy for an hour at a low laser setting but its adhesions dimmed, by 15 % in brightness and 24 % in thresholded area, and at 8– 10 % the fluorescence halved within 25–50 s of exposure and the cells retracted (Figure 4; Figure S4). The continuous-wave recordings were made on a commercial microscope optimised for that purpose, whereas continuous-wave lines coupled into the two-polarisation path of the pulsed instrument gave poor images; with respect to the quality of illumination the comparison therefore favours the continuous-wave case. What it does not control is the specimen (cell line and labelled protein) and the irradiance, which cannot be matched between a 20-nm band and a laser line and was set on each instrument by image contrast at the camera (Limitations of the study).

### Relation to other single-molecule approaches

The single-molecule part of this work is closest to single-molecule speckle microscopy, in which individual fluorescent-protein fusions are seen as spots while they are immobilised on a structure, and freely diffusing molecules blur into the background^19,20^; the approach has been applied to actin flow over adhesions and to talin itself^16,21^. Speckle microscopy obtains sparsity from a very low density of labelled molecules. Here talin is expressed at ordinary levels, and sparsity arises because only a small fraction of the bound GFP molecules emit at any instant, as in localisation methods that exploit the blinking of conventional fluorescent proteins^22,23^. In contrast to single-particle-tracking PALM of adhesion proteins^24,25^, there is no photoactivation step and molecules are not followed individually: what is measured is the slow collective translation of the immobilised population over tens of minutes, a time scale complementary to the tens of milliseconds of single-particle tracks. Localisation counts are therefore detection events, not numbers of molecules.

### Photobleaching in ordinary TIRF imaging: what the schedule contributes

Single-molecule detection is the most demanding use of the illumination described here, but it is not the only one, and framing the result as an SMLM method understates where it is most immediately useful. The great majority of live-cell TIRF experiments are diffraction-limited: a labelled structure is followed over minutes to hours, and the experimenter trades frame rate, exposure and laser power against how long the signal lasts. In that trade the limiting quantity is the same photochemistry, and the long-duration dataset reported here is an ordinary time-lapse experiment, not a single-molecule one — 100–200 ms exposures at 1–10 s intervals, with adhesions resolved as diffraction-limited structures. Its outcome is that a full hour at one frame per second can leave the adhesion fluorescence at or above its starting value (1.03 and 1.37; a third hour-long recording ended at 0.54), and that an eight-hour recording at a 2 % duty cycle retained about half (0.46). The continuous-wave recordings place this result. At the same schedule, continuous-wave TIRF on a commercial microscope also sustained an hour at one frame per second at a 2 % laser setting, with the cell footprint and adhesion motility unchanged, so for diffraction-limited imaging of a bright structure a low frame-level duty cycle and a low laser setting go most of the way on their own. What differed was the read-out of the fluorophore: adhesion brightness of 0.85 and thresholded adhesion area of 0.76 under continuous-wave excitation, against 1.03 and 1.05 under pulsed excitation, with the adhesions of the continuous-wave cell fading at constant size and those of the pulsed cell growing (Table 1). At 8–10 % the same schedule halved the footprint fluorescence within 25–50 s of exposure and the cells retracted, at cumulative exposures at which every hour-long pulsed series of this study was still at or above its starting signal.

Two features of the schedule are separable, and they carry different practical weight. The frame-level duty cycle — illuminating for a fraction of each interval — is available on every microscope with a shutter or an acousto-optic gate, costs nothing, and is widely used; the only claim we make about it is quantitative, namely how far it can be pushed with this source before the signal decays. The pulse-level duty cycle, which caps the rate at which any individual molecule can be driven round the excitation cycle at 10⁶ s⁻¹ regardless of local intensity, requires a pulsed source and a pulse picker, and it is the part of the recipe that has no counterpart in a continuous-wave instrument. Every pulsed series was recorded at 1 MHz, so the present data do not separate their contributions (see the next section). The usefulness of the frame-level result does not depend on that separation: an hour of pulsed TIRF with no net loss of GFP signal is a useful acquisition recipe whether the mechanism is pulse-level photophysics, the frame-level recovery interval, or both.

A second general point concerns how photobleaching is reported. Every declining trace in a live-cell experiment bounds photobleaching from above and is not a measurement of it, because focus and TIRF-angle drift, cell migration out of the evanescent field, protein turnover and stress responses all reduce the signal and none of them are reversible by the fluorophore. A flat or rising trace, by contrast, cannot be produced by any of those confounders, since nothing restores a bleached fluorophore. We therefore treat the non-declining recordings as the informative ones and the declining recordings as uninformative, and we report values per recording rather than a population mean; the dual-colour acquisitions make the same argument internally, since paxillin-mApple fell in every recording while GFP-talin rose in several of the same recordings, under identical illumination, focus and cell state. That internal control is weaker than it would be with two labels on one protein, because talin and paxillin differ in turnover as well as in fluorophore, and it is reported as such.

### Why pulsed excitation might preserve GFP: the photophysics

The published photophysics offers several candidate explanations for the difference between the pulsed and the continuous-wave hour, and does not yet single one out.

#### Relaxation of dark states between pulses

Donnert et al.^5^ found that separating the excitation pulses by more than 1 µs, instead of 25 ns, increased the total fluorescence obtained from GFP before bleaching by up to about 25-fold, for one- and two-photon excitation, and concluded that a dark state with a lifetime of about a microsecond is the precursor of bleaching. Our 1-µs interval satisfies that condition. Two qualifications apply. The dark state concerned cannot be the lowest triplet state of the chromophore as now characterised, which lives for about 5 ms^7^; the earlier estimates of 2–25 µs for the triplet lifetime of GFP variants (quoted by Boudreau et al.^8^) preceded this direct measurement, so in GFP the microsecond state is either a different species or the gain has another origin. And the gain of Donnert et al. was obtained at the irradiance of a focused beam, where a molecule under continuous-wave light is re-excited within much less than a microsecond. In wide-field TIRF the continuous-wave excitation rate of EGFP (absorption cross-section ≈ 2 × 10⁻¹⁶ cm² at 488 nm, from an extinction coefficient of 56,000 M⁻¹ cm⁻¹) is about 5 × 10⁵ s⁻¹ at 1 kW cm⁻² and 5 × 10³ s⁻¹ at 10 W cm⁻². At single-molecule irradiances about 40 % of successive absorptions are therefore less than 1 µs apart, and pulse picking removes them; at the irradiances of ordinary time-lapse fewer than 1 % are, and pulse picking changes little. Our recordings were made at an estimated 1–3 × 10⁴ excitations per molecule per second during an exposure (Methods), at which continuous-wave light would place only 1–3 % of successive absorptions less than 1 µs apart. If this estimate holds, relaxation of a microsecond dark state between pulses can account for little of the difference, in the single-molecule recordings as in the hour-long time-lapse.

#### The triplet state

With a formation yield of about 1 % and a lifetime of about 5 ms, the triplet state of EGFP holds half of the molecules at an excitation rate of 2 × 10⁴ s⁻¹, within the range estimated for our recordings, and approaches that level within a few milliseconds of the start of an exposure; it absorbs throughout the visible and near-infrared and is the gateway to oxidative photochemistry^7^, and depopulating it with near-infrared light reduces the bleaching of fluorescent proteins 1.5–9-fold^11^. Its population depends on the average excitation rate and not on whether the light arrives as a 1-MHz pulse train, because thousands of pulses fall within one triplet lifetime. What does give the triplet state time to empty is a dark interval of milliseconds or more: the 80 % or more of each frame period without illumination in the schedules used here, the return time of a scanning beam^9^, or the line averaging with which Boudreau et al.^8^ reduced the bleaching of EGFP nine-fold. This part of the recipe is shared with our continuous-wave recordings. A large standing population of dark molecules is also what the single-molecule data show: most bound GFP-talin was dark at any instant, and the same sites emitted again and again over 30 min (Note S1).

#### Irradiance

Photobleaching of fluorescent proteins is supra-linear in irradiance, with exponents of 1.07 for mEGFP in vitro and about 1.2–1.3 in cells^10^, and our continuous-wave series shows the same behaviour in an extreme form. Any scheme that forms its image at a lower irradiance therefore gains disproportionately. The windowed supercontinuum delivered little average power — too little to register on a thermal sensor, an estimated 60 W cm⁻² during an exposure (Methods), and twenty-fold less after pulse picking than the unpicked pulse train — and the EMCCD camera formed usable images from it (single-frame signal-to-noise ratio 10, against 56 in the continuous-wave recording). A low average irradiance may thus be part of the explanation of the pulsed result, and to that extent the benefit would also be available to a continuous-wave source attenuated to the same level and read out with the same camera. Conversely, the peak irradiance within a picosecond pulse is 10⁴–10⁵ times the average, and to the extent that the supra-linearity reflects absorption by short-lived excited states^2^, pulsing would be expected to increase bleaching. Boudreau et al.^8^ nevertheless found that 250-ps pulses at 20–80 MHz halved the bleaching of EGFP relative to continuous illumination at equal exposure, with no dependence on the repetition rate — an observation that none of the mechanisms above explains, and the closest published precedent for a benefit of picosecond pulses as such.

#### Homogeneity of the field

Two further properties of the source are unrelated to pulsing but may contribute. The 20-nm bandwidth and the two time-separated, orthogonally polarised beams give an evanescent field free of the interference fringes of single-mode laser TIRF, so that no part of the cell is exposed to more than the mean irradiance — an advantage that grows with the supra-linearity of bleaching — and the excitation is independent of the in-plane orientation of the fluorophore. A homogeneous field also lets the irradiance be set by what the dimmest part of the cell needs: with a fringe contrast m, the mean must rise by 1/(1 − m) for the minima to reach a given level, twofold for m = 0.5. This argument remains qualitative until the uniformity of both fields is measured on a uniform fluorescent layer.

Separating these contributions requires one specimen under both sources at measured irradiance, with the repetition rate varied (Outlook).

### Cost and availability of the illumination

For adoption, the limitation that weighs most is the source. The supercontinuum laser used here delivers 8 W spread over about 2000 nm, of which a 20-nm band carries of the order of 1 %, and it needed a four-channel acousto-optic tunable filter and a second TIRF illuminator to use that band efficiently (Methods). What the pulsed recordings require is much less: pulses far shorter than the excited-state lifetime, an estimated 1.6 nJ per pulse in a 50-µm field (Methods, equations 1–3), at a repetition rate of 1 MHz — an average power of about 1.6 mW. The requirement is therefore a pulse energy at a low repetition rate, not a high average power. Table 2 compares the source used here with two kinds of source that meet this specification at lower cost, and with the continuous-wave line of the comparison instrument. Gain-switched picosecond diode lasers set their repetition rate electronically, from single shots to tens of megahertz, so that no pulse picker is needed, and high-power heads at 470–485 nm are specified at pulse energies of up to about 2 nJ^26^; supercontinuum lasers of lower total power than ours, with pulse pickers, are sold for fluorescence-lifetime and STED microscopy^27^. Both kinds of source are installed in many imaging facilities, where the pulsed excitation described here could be tried on an existing TIRF stand. None of these alternatives was tested in this work, and the prices in Table 2 indicate orders of magnitude only.

**Table 2.** Excitation sources for low-duty-cycle pulsed TIRF: the source used here, lower-cost alternatives and the continuous-wave reference.

| **Source** | **Pulse duration and repetition rate** | **Energy per pulse** | **Further hardware for TIRF** | **Indicative price, excitation hardware only** |
|---|---|---|---|---|
| This work: supercontinuum laser with pulse picker, 8 W (Fianium WL-SC450-8) | 20–80 ps; 20 MHz picked to 1 MHz | ≈ 4 nJ in a 20-nm band at the source, ≈ 1.6 nJ in the field (estimate; Methods) | four-channel acousto-optic tunable filter; polarisation splitting into two fibres; second TIRF illuminator | ≈ €100,000–150,000 |
| Supercontinuum laser of lower total power with pulse picker (as used for FLIM and STED) | picoseconds; picked from the fundamental, e.g. 2–78 MHz, lower on request | total power divided by the fundamental repetition rate, spread over the whole spectrum (400 nJ for the source used here) | acousto-optic tunable filter or band-pass filter; polariser or polarisation splitting | ≈ €40,000–80,000 |
| Picosecond pulsed diode laser, 470–485 nm, high-power head | < 100–220 ps; set electronically, single shot to 50 MHz | up to ≈ 2 nJ at the laser head | driver; fibre coupling into a standard TIRF illuminator; linearly polarised, so no splitting | ≈ €15,000–30,000 |
| Reference: continuous-wave 488 nm laser line, 60 mW (Omicron LightHUB+; comparison instrument) | continuous | — | none beyond a standard TIRF illuminator | ≈ €10,000–20,000 per line |

Prices are indicative 2026 list prices of the excitation hardware alone, excluding microscope, TIRF illuminator, camera and taxes, and vary with configuration and supplier. Pulse energies of the alternatives are manufacturers’ specifications^26,27^; the 470-nm heads excite GFP less efficiently than 488 nm. None of the alternatives was tested in this work.

### Outlook

The decisive experiment is simple to state: one TIRF microscope with a picosecond source and pulse picker and a continuous-wave laser at the same wavelength, both measured at the sample; an immobilised or fixed GFP specimen in which exchange is absent; and a repetition-rate series from 20 MHz to 100 kHz at constant average power. Supercontinuum and picosecond diode lasers with pulse pickers are standard equipment in fluorescence-lifetime and STED laboratories, so the experiment is within reach of many groups, and the acquisition parameters, data and analysis tools reported here are intended to make our side of the comparison reproducible. Independently of its outcome, the recordings support three practical points for live-cell TIRF of GFP fusions. First, a frame-level duty cycle of 10–20 % at a low laser setting is compatible with an hour of imaging at one frame per second on both instruments, with cell footprint and adhesion motility unchanged. Second, under continuous-wave excitation the laser setting, not the cumulative exposure, limited the experiment: raising the setting five-fold shortened the usable exposure by more than an order of magnitude, and brought cell retraction with it. Third, bleaching in living cells is best reported per recording, with a read-out that tracks the labelled structure together with independent measures of cell behaviour, because a declining trace alone does not distinguish photobleaching from biology.

### Limitations of the study

Four limitations are evident. First, the source used here is expensive: a high-power supercontinuum laser with a pulse picker and a multichannel acousto-optic tunable filter costs considerably more than the continuous-wave lasers of a standard TIRF microscope. The pulse condition itself does not require such a source, and lower-cost sources that meet it are common in fluorescence-lifetime and STED facilities (Table 2), but none of them was tested here. Second, the gain is smaller for red fluorescent proteins. mCherry bleaches considerably faster than GFP in general, and did so in the same cells here (Notes S1 and S3); paxillin-mApple likewise ended below its starting brightness in every dual-colour recording of the long-duration dataset (Note S4). Under pulsed excitation with the 561-nm band of the same source, mCherry nevertheless bleached more slowly than we observe under continuous-wave TIRF, so for an end user imaging red fusions the method is an improvement, but it may not be enough for recordings as long as those reported here for GFP. A likely reason for the smaller gain is that bringing mCherry into its triplet state to the extent estimated here for GFP (Discussion) requires substantially more excitation power than the 561-nm band of our source could deliver; pulsed excitation of red fluorescent proteins at higher power is an interesting follow-up experiment. Third, the irradiance at the sample was measured for neither source and cannot be matched by setting alone between a 20-nm band and a laser line; illumination was set by image contrast at the camera, and a calibrated comparison needs a power measurement at the objective for both sources. Fourth, talin exchanges with a cytosolic pool that TIRF does not illuminate, so retained fluorescence reflects the survival of fluorophores together with their replenishment; a fixed or non-exchanging specimen would separate the two. We no longer have direct access to the pulsed instrument, so the last three tests fall to instruments with comparable sources. The work demonstrates a method rather than a biological effect; the pulsed and continuous-wave hours were recorded on different microscopes, the schedule-matched comparison rests on one hour-long recording of each, and the number of recordings per condition is small; all per-recording values are reported (Tables S1–S4; Figure S5).

## Methods

Cell lines, constructs, reagents, instruments and software are listed in Table S5 (Supplementary Material).

### Cells and constructs

#### Cell lines

The single-molecule series (2017) were recorded in MDA-MB-231 human breast cancer cells (female) expressing GFP-talin-mCherry, i.e. talin with GFP fused to the N-terminal head domain and mCherry to the C-terminal rod, the construct described by Margadant et al.^28^ and Hu et al.^29^. All localisation analyses use the GFP channel; the mCherry channel is used only for the ensemble bleaching comparison and the co-detection test. The long-duration pulsed series (2018) were recorded in MDA-MB-231 human breast cancer cells (female) co-expressing GFP-talin and paxillin-mApple as two separate proteins. The continuous-wave series (2026) were recorded in U2OS human osteosarcoma cells (female) expressing vinculin-GFP. MDA-MB-231 and U2OS cells and the GFP-talin, paxillin-mApple and vinculin-GFP plasmids were obtained from the laboratory of Michael P. Sheetz (Mechanobiology Institute, National University of Singapore); the GFP-talin-mCherry plasmid was that of Margadant et al.^28^ and Hu et al.^29^. Cells were transfected with jetOPTIMUS DNA transfection reagent (Polyplus, Sartorius; Cat# 201000001) according to the manufacturer’s protocol 24 h before imaging and were imaged in 35-mm glass-bottom dishes (MatTek P35G-1.5-14-C or ibidi µ-Dish 35 mm, high, Glass Bottom, 81158). Cells were cultured and imaged in DMEM supplemented with 10 % fetal bovine serum; during imaging they were kept in an on-stage incubator at 37 °C and 5 % CO₂. The cell lines were tested regularly for mycoplasma contamination.

### Instrument, acquisition and analysis

#### Pulsed supercontinuum TIRF microscope

The pulsed recordings were made on a Nikon Ti-E with N-STORM TIRF illuminator, HP Apo TIRF 100×/1.49 objective, Perfect Focus active, and a Photometrics Evolve Delta EMCCD (EM gain 400, 20 MHz readout, conversion-gain setting 2, −80 °C). With the Ti-E 1.5× intermediate magnification lens (cells 1 and 2, and the GFP-channel stream) the pixel size is 107 nm; without it (cell 3) 160 nm. The acquisition software recorded 160 nm in all cases; the correct value was applied in all analyses.

#### Excitation source

The source was a picosecond supercontinuum laser (Fianium WL-SC450-8, 8 W total average output, fundamental repetition rate 20 MHz, internal pulse picker) coupled into the N-STORM illuminator (Figure 1a, d; Figure S3). The supercontinuum output was spectrally windowed by a four-channel acousto-optic tunable filter (each channel rated for 2 W); all four channels were set to the same 20-nm band (centred at 488 nm or 561 nm) to recover the weak in-band power and to minimise spectral cross-talk, and their outputs were combined into a single multimode fibre. The transmission setting was 100 % in all recordings. Pulse duration at the fibre output is **5–20 ps in the low-power mode and 20–80 ps in the high-power mode** used here. This is an intermediate pulse-duration regime: supercontinuum generation in the fibre broadens the single-digit-femtosecond pulses of the seed oscillator both spectrally, across 400–2200 nm, and temporally, by some four orders of magnitude. The distinction matters because peak irradiance at a given average power scales inversely with pulse duration, so picosecond pulses deliver roughly 10³–10⁴ times lower peak irradiance than femtosecond pulses of the same average power, keeping the excitation in the linear single-photon regime.

#### Two-module, polarisation-split TIRF illumination path

The supercontinuum is unpolarised, which is incompatible with a single-beam TIRF illuminator that requires a fixed polarisation relative to the plane of incidence (Figure 1b, d). Rather than discard half of the power with a polariser, the light was split into its two orthogonal linear polarisation components, which were delivered by separate fibres to two Nikon TIRF illuminators mounted on the two layers of the Ti-E filter turret (recorded in the acquisition metadata as the N-STORM port of the lower layer and port 2 of the upper layer). The two fibres differ in length by 3 cm, so the pulses of the two polarisations arrive about 150 ps apart; they do not overlap in time (pulse duration 20–80 ps), but both fall well within the excited-state lifetime of the fluorophore. The two illuminators deliver the beams to the objective back focal plane at azimuths 90° apart, each with its polarisation set with respect to its own plane of incidence, at angles of incidence of 63.6–66.6° (critical angle 62.7°); the angle was set for each illuminator separately and was not always the same in the two, so the penetration depths of the two components could differ somewhat. Three properties of this path matter for single-molecule imaging: (i) the full in-band power is used; (ii) excitation from two orthogonal azimuths with two polarisations is independent of the in-plane orientation of the fluorophore; and (iii) the 20-nm bandwidth and the time-separated beams give low coherence, so the TIRF field is free of the interference fringes and scatter patterns typical of single-mode laser TIRF — the illumination resembles that of an LED, which is what the raw frames show. All series reported here — the two-channel series of cells 1–3 and the GFP-channel stream — were recorded with the pulse picker set to 20:1, i.e. at 1 MHz with 1 µs between pulses, following the advice not to cycle GFP above ∼1.2 MHz. The picked rate was verified on a fast photodiode and oscilloscope, whose frequency counter read 984.937 kHz (Figure S3f); the pulse duration is below the bandwidth of that instrument and is taken from the manufacturer’s specification. The pulses (picoseconds) are much shorter than the GFP excited-state lifetime (∼2.5 ns), and the 1 µs interval is some 400 times that lifetime, so a molecule excited by one pulse has returned to the ground state, or has crossed to a dark state, long before the next pulse arrives. The interval is three orders of magnitude shorter than the lifetime of the EGFP triplet state^7^ (about 5 ms), so pulse picking does not by itself let the triplet state empty between pulses; it does correspond to the pulse separation (> 1 µs) above which Donnert et al.^5^ observed an increased fluorescence yield from GFP. Because each pulse is much shorter than the excited-state lifetime, a molecule completes at most one excitation cycle per pulse, and what pulse picking sets is a ceiling on how fast any individual molecule can be cycled — at most 10⁶ excitations per second, irrespective of local intensity — and, at fixed pulse energy, an average power twenty-fold below that of the unpicked 20 MHz train (Discussion). TIRF angle 63.6–66.6° (critical angle 62.7°).

#### Excitation power

The average power of the windowed bands could not be measured with the sensors available: a silicon photodiode head saturates on the picosecond peaks, and a calorimetric head was below its sensitivity floor at the few-milliwatt average level. At the 1 MHz rate used for every series reported here, a 20–80 ps pulse gives a pulse-level illumination duty cycle of 1:50,000 to 1:12,500, so the peak irradiance exceeds the average by the same factors (it would be 1:2,500 to 1:625 at the unpicked 20 MHz). The frame-level duty cycle of the acquisition is separate from this: a 200 ms exposure every 2.07 s, i.e. 1:10 (Figure 1c). The excitation probability per pulse was instead estimated from the specification of the source. If the 8 W average output at 20 MHz (400 nJ per pulse) is spread evenly over the ≈ 2000 nm of the supercontinuum, the spectral energy density of one pulse is dE/dλ ≈ 0.2 nJ nm⁻¹; pulse picking selects pulses without changing their energy. The number of photons per pulse in a band from λ₀ to λ₀ + Δ that reach the field of view, when a fraction η of the in-band energy arrives there, is

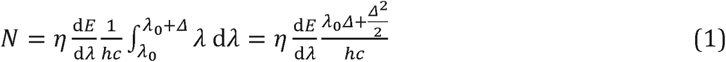

and the photon fluence per pulse over a square field of side a is

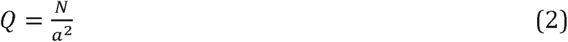

A molecule of absorption cross-section σ in a field of area A that receives QA photons escapes excitation with probability (1 − σ/A) raised to the power QA, so the probability that it is excited by one pulse is

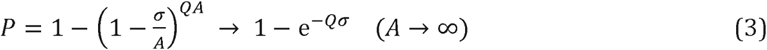

With η = 0.4, a = 50 µm, Δ = 20 nm and λ₀ = 478 nm (the 488-nm band), N ≈ 4.0 × 10⁹ photons and Q ≈ 1.6 × 10¹⁸ m⁻² (1.6 × 10¹⁴ cm⁻²) per pulse. For σ between 6.5 × 10⁻¹⁷ cm² and 2.1 × 10⁻¹⁶ cm² (the latter for EGFP at its absorption maximum, from ε = 56,000 M⁻¹ cm⁻¹), Qσ = 0.010–0.034 and P ≈ 1–3 % per pulse. At 1 MHz this corresponds to 1–3 × 10⁴ excitations per molecule per second during an exposure and to an average irradiance of about 60 W cm⁻² (1.6 mW over the field). Because the spectral density of a supercontinuum is not uniform and the transmitted fraction was not measured, these are order-of-magnitude estimates; they place the excitation far from saturation, so the fluorescence is expected to scale linearly with pulse energy.

Because the excitation spectra of the two sources differ (a 20-nm band against a laser line) and neither power was measured, illumination was set on each instrument at the level of the camera read-out, to give comparable image contrast of the specimen; the single-frame signal-to-noise ratios actually obtained are given in Table 1. Continuous-wave 488 nm and 561 nm lasers on the same microscope gave single frames of comparable brightness at the settings used, but no CW time series were recorded for this dataset; the continuous-wave power series of Figure 4 was recorded in 2026 on a different instrument (below).

#### Acquisition of the single-molecule series

Two-channel series: at each time point the GFP channel (488-nm band, 200 ms exposure) and the mCherry channel (561-nm band, 300 ms) were acquired sequentially; time-point interval 2.07 s (cell 1, 283 time points), 2.10 s (cell 2, 901) and 2.08 s (cell 3, 901). GFP-channel stream: 1818 frames of 330 ms with no interval (0.33 s per frame, 100 % duty cycle, 10.0 min). All recordings were made on 2017-05-19 at 37 °C.

#### Long-duration pulsed series

MDA-MB-231 human breast cancer cells co-expressing GFP-talin and paxillin-mApple (two separate proteins) were imaged on the same microscope and camera on 2018-08-03, -05 and - 06 at 160 nm per pixel with no intermediate magnification (N-STORM Zoom = 1x in the acquisition metadata, so the recorded calibration is correct for these series, unlike cells 1 and 2 above), EM gain 185–300, TIRF angle 68–75°. The per-line AOTF settings recorded in the acquisition metadata differ between the three hour-long series held as raw ND2 files: the series that ended at 0.54 of its starting adhesion brightness lists all four 488-nm AOTF lines at 100 %, the series that ended at 1.37 lists line 1 at 100 % and lines 2–4 at 0 %, and the series of Figure 3 lists line 1 at 50 % and lines 2–4 at 0 % (EM gain 300, 20 MHz readout); how these settings map onto power at the sample was not determined. Applied to the raw file of the series of Figure 3, the adhesion-brightness procedure below returned an end-to-start ratio of 1.03 (tabulated 1.03), and phase correlation of 10-frame means at 5-min intervals gave a drift below 1 pixel over the hour. Nineteen acquisitions were analysed: a 100 ms exposure (200 ms in two series) at a fixed period of 1, 2, 5 or 10 s, 132–5761 frames, 4.4 min to 8.0 h; ten acquisitions recorded both channels in alternation within each time point and nine the GFP channel only (Table S3). For each acquisition a cell mask was obtained from the smoothed time-average of the channel and a background region from its complement dilated by 12 pixels; the background was taken per frame as the median over that region. Two read-outs were computed per frame: the mean of the cell mask minus the background, and the adhesion brightness, defined as the 99th percentile of the frame minus the same background. The adhesion brightness is the quantity reported in Results and Figure 3 because the fixed-mask mean is inflated when a cell spreads into the mask and deflated when it retracts, whereas the upper percentile tracks the labelled structures themselves; both are tabulated in Table S3 and agree to within 0.1 in 23 of the 29 cell-channels, the largest discrepancy being 0.33 (one GFP channel: 1.48 by fixed-mask mean, 1.82 by adhesion brightness). End-to-start ratios are the mean of the last 1 % of frames divided by the mean of the first 1 %.

#### Morphometry and adhesion tracking of the hour-long pulsed recording

All measurements for Figure 3 were made on the raw ND2 file at 160 nm per pixel without drift correction. Adhesions were segmented every 10 frames on the mean of five consecutive frames: the image was smoothed (Gaussian, σ = 1 pixel), a local background (Gaussian, σ = 8 pixels) was subtracted, and pixels more than 5 times the robust noise of a cell-free region (7.2 counts) were retained if they belonged to objects of at least 4 pixels (0.10 µm²) within 10 pixels of the cell footprint; one fixed threshold was used for the whole hour. The cell footprint was the region in which the 30-frame mean, smoothed with σ = 4 pixels, exceeded the per-frame background of a cell-free region by 7 counts (5 and 10 counts for the band in Figure 3d), after morphological closing, hole filling and removal of objects below 50 µm²; subtracting the per-frame rather than a fixed background is necessary because the camera background fell by 7 counts over the hour, comparable to the diffuse signal. Adhesion centroids (weighted by background-subtracted intensity) were linked between successive time points by minimum-cost assignment with a maximum displacement of 3 pixels (0.48 µm) per 10 s; tracks of at least 18 time points (3 min) were kept, their speed taken as the slope of a linear fit of position against time, and their direction relative to the centroid of the footprint. The noise threshold for movement was the 95th percentile of the same fits after random permutation of the time points within each track. Centroid displacement includes changes of adhesion shape as well as translocation, so the speeds are those of the adhesion centroid, not of the molecules within it.

No drift correction, frame exclusion or cropping was applied to these series, since the read-outs are intensity statistics over masks rather than positions. Timing was taken from the Equidistant (Period …) field of the per-acquisition metadata export where present (14 of 19) and otherwise from the ND2 plane timestamps; the Time [s] field is written in milliseconds in some files of this dataset and in seconds in others, and the interpretation giving an interval in the 0.05–120 s range was used. One acquisition has neither a metadata export nor recoverable timestamps and is tabulated without a time axis. Whole-field means were not used as a read-out: they include a background pedestal that is 11–91 % of the total signal (median 41 %), which compresses the apparent fractional change by the same factor.

#### Continuous-wave power series

U2OS cells expressing vinculin-GFP were imaged on 2026-11-06, 09, 10 and 11 on a Nikon Ti2-E with the H-TIRF module, Apo TIRF 100×/1.49 oil objective and Photometrics Prime 95B sCMOS camera (200 MHz readout, sensitivity mode). Excitation came from the 488 nm line of a continuous-wave laser combiner (Omicron-Laserage LightHUB+, MOM50201 LH+APC; 405/488 nm, 60/60 mW; maximum output 60 mW at 488 nm), set to 1 % (one recording), 2 % (one), 5 % (two), 8 % (two) or 10 % (three), as recorded in the acquisition metadata, which list no other active line; no brightness trace showed a step change. These settings correspond to nominal outputs of 0.6–6 mW at the laser, before fibre coupling and the TIRF illuminator; the power at the objective was not measured at any setting. The H-TIRF angle recorded in the metadata was 65.3° for the 1 % recording (nominal penetration depth 118 nm), 68.7° for the 2 % recording (99 nm), 72.6° for the 5 % recordings (77 nm) and 67.6° for the 8 % and 10 % recordings (110 nm). Pixel size was 110 nm, except for four of the 8–10 % recordings, which were recorded with an additional 1.5× magnification (73 nm per pixel); this changes the signal per pixel but not the irradiance at the sample, and all read-outs are normalised to the start of each recording. Exposures of 100 ms were taken every 1.0 s, except for the 1 % recording (streamed at 0.117 s per frame); the second 5 % recording is a continuation of the field of the first, cropped to 804 × 804 pixels. Adhesion brightness was computed exactly as for the 2018 series (cell mask by Otsu thresholding of the smoothed time-average, background as the per-frame median of the region outside the mask dilated by 12 pixels, 99th percentile of the frame minus that background, end-to-start ratio of the means of the last and first 1 % of frames); applied to the two 2018 series held as raw ND2 files, this implementation returned 0.53 and 1.38 against the tabulated 0.54 and 1.37. Footprint fluorescence was the background-subtracted sum over the initial cell footprint (threshold at 8 % of the 5th–99.5th percentile range of the smoothed mean of frames 2–11, largest component, filled, dilated by 1 µm; background from the region more than 5 µm outside the footprint and below the 30th percentile), normalised to the first 1 % of frames; for the pulsed comparison the fixed-mask mean of Table S3 was used. Exposures to 0.8 and 0.5 were the first crossings of a 9-frame running median, and the retention at 45 s (Figure 4e) the value of a 9-frame running median at that cumulative exposure. Footprint area at 1–5 % was measured per frame at 15 % of that frame’s own median footprint intensity above background, so that uniform dimming does not register as retraction. At 8–10 %, where most of the signal was lost, this threshold admits background, and area was instead measured on 2× binned frames by Otsu thresholding of the log-transformed, smoothed (σ = 2 pixels), background-offset frame, followed by morphological opening, hole filling and retention of objects ≥ 150 µm² overlapping the first-frame mask dilated by 3 µm; masks were inspected on first and last frames, and the result is reported only for the two isolated 8 % cells, for which they followed the cell edge throughout. The single-frame adhesion signal-to-noise ratio was the mean background-subtracted intensity of the brightest 5 % of cell pixels in frames 2–21 divided by the temporal standard deviation of background pixels over the same frames; it is camera-independent only to first order and is used to indicate the imaging regime, not to match excitation. Per-frame read-outs, sub-sampled 2–4× binned image stacks and movies of every series are archived with the source data.

#### Schedule-matched continuous-wave recording

The hour-long 2 % recording (Figure S4; Table 1) was analysed with the procedure used for the pulsed recording of Figure 3, with all lengths converted from 160 nm to 110 nm pixels: smoothing σ = 0.16 µm, local-background σ = 1.28 µm, minimum adhesion size 0.10 µm², footprint smoothing σ = 0.64 µm, maximum linking distance 0.48 µm per 10 s, tracks of at least 3 min. The noise of the sCMOS images is far lower than that of the EMCCD images of the pulsed recording (robust noise of the high-pass image 0.8 against 7.2 counts), so a threshold defined as a multiple of the noise would segment diffuse texture; the adhesion threshold was therefore set at the same fraction of the initial adhesion brightness as for the pulsed recording (0.158, i.e. 20.3 counts for an initial adhesion brightness of 128.7 counts; for the pulsed recording, 35.9 counts for 227 counts), and one fixed threshold was used for the whole hour. Because cells fill most of the field, the background was taken per time point as the median of the pixels below the 12th percentile of the smoothed time-average; the cell footprint was the region in which the smoothed 30-frame mean exceeded this background by 4 counts (3 and 6 counts for the range in Table 1), after morphological closing, hole filling and removal of objects below 50 µm². Masks were inspected on the first and last frames. Tracking gave 1971 tracks of at least 3 min (median 4.7 min); the 95th percentile of the time-shuffled speeds was 89 nm min⁻¹, against 32 nm min⁻¹ for the pulsed recording, reflecting the higher density of adhesions and the correspondingly larger centroid noise, so absolute speeds are not compared between the two recordings. The raw ND2 file (18.6 GB) was converted to a losslessly compressed OME-TIFF cropped to columns 0– 1500 of 1608 (7.6 GB); each of the 3600 frames was verified against the raw data by SHA-1 hash before the raw file was removed, and the complete ND2 metadata are stored alongside.

#### Data handling

ND2 files were converted losslessly to OME-TIFF with the complete acquisition metadata embedded (Python nd2 0.11, tifffile; verification by byte-wise comparison of every plane). Nine of the nineteen 2018 acquisitions are held as cropped big-endian TIFF exports carrying an .nd2 extension; these were read as TIFF and are flagged in the file manifest (Data S1), and the corresponding raw ND2 files are not present in that dataset.

#### Pre-processing

Frames whose mean intensity deviated from a 21-frame rolling median by more than five robust standard deviations (transient intensity spikes or dips) were discarded (27 of 901 frames in cell 2; 29 of 1818 in the stream; none in cells 1 and 3); the field was cropped to the padded bounding box of the cell where stated.

#### Drift correction

Ten-frame (cells 1 and 3), twenty-frame (cell 2) or thirty-frame (stream) averages of the GFP channel were registered to the first average by phase cross-correlation with 20-fold upsampling (scikit-image); the trajectory was smoothed with a cubic smoothing spline and applied to each frame (and to the mCherry channel of the same time point) by bilinear interpolation. Residual drift was assessed by cross-correlating localisation histograms (2× camera-pixel binning) of consecutive 25-frame segments. The chromatic offset between channels was measured by cross-correlation of the time-integrated localisation maps and applied to the mCherry coordinates before the co-detection analysis.

#### Localisation

Each frame was band-pass filtered with a difference of Gaussians (σ = 107 and 268 nm, i.e. 1.0 and 2.5 pixels at 107 nm per pixel); local maxima exceeding the median plus four robust standard deviations (1.4826 × MAD) within a 7 × 7 neighbourhood (5 × 5 at 160 nm per pixel) were fitted by least squares with a symmetric two-dimensional Gaussian plus constant background in a window of 0.96–1.1 µm (scipy.optimize.least_squares). Fits with σ outside 86–235 nm or non-positive amplitude were rejected. Intensities are reported in camera units because the detector’s electron-per-count conversion at the gain setting used was not determined.

#### Linking and event statistics

Detections in consecutive frames within 150 nm were linked into tracks (nearest neighbour, no gap closing); the fraction of single-frame tracks, the track-duration distribution and the next-frame persistence (fraction of detections with a successor) were computed per series. The mean bright-state duration was estimated from the persistence p at frame interval Δt as Δt/(1 − p) under the assumption of a constant hazard. Site recurrence was quantified as the number of detection pairs within 40 nm and their time lags, compared with the same count after displacing one copy of the coordinates by 250 nm. The fraction of adhesion fluorescence accounted for by detected events was computed per frame as the sum of fitted spot integrals within the adhesion mask divided by the background-subtracted integral of the raw frame over the same mask.

#### Adhesion analysis in the localisation data

Localisations were binned at 50 nm, smoothed (σ = 1.5 bins) and thresholded at the 90th percentile of non-zero density; connected regions defined adhesions. For the flow analysis, adhesions with ≥ 150 localisations spanning ≥ 6 min were used; the principal axis was obtained from the covariance of the localisation coordinates and oriented toward the cell centre (mean of all localisations), and the Theil–Sen slope of projected position versus time gave the along-axis speed, with significance from the ordinary least-squares regression p-value. For the centroid analysis, adhesions with ≥ 40 localisations in both the first and the last third of the series were used. Wilcoxon signed-rank tests were two-sided.

#### Software

Analyses were run in Python 3.13 with nd2 0.11.3, tifffile 2026.6.1, scikit-image 0.26, SciPy, NumPy and Matplotlib. ImageJ plugins implementing the format conversion, drift correction and localisation (Java/Groovy) and the Python analysis code (PulsedTIRF_analysis), which reproduces the source data of every figure and table, are provided (data and code availability).

### Quantification and statistical analysis

Each recording is one cell or one field of view from one dish. The single-molecule dataset comprises three 2-s series, six 5-s series and one streamed series, all recorded on 2017-05-19; the long-duration pulsed dataset comprises nineteen acquisitions from three days in 2018-08, three of which lasted a full hour; the continuous-wave dataset comprises nine recordings from four days in 2026-11. No recordings were excluded; frames excluded for transient intensity spikes are stated per series. Because the study demonstrates a method and the numbers of recordings are small, values are reported per recording and no population statistics are computed across cells; tests within a recording (Wilcoxon signed-rank, regression slopes, permutation thresholds) are two-sided and are described with the corresponding analysis. Numbers of adhesions and tracks entering each analysis are given in the figure legends and in Tables S1–S4. The recording shown in Figure 3 was selected for display from the three hour-long pulsed recordings, all of which are shown in Figure S5.

## Supplementary Material

Supporting Figures S1–S5, Tables S1 and S5 and Notes S1–S5 follow the References in this file; Tables S2–S4, Videos S1–S2 and Data S1 are provided as supplementary files.

- Table S2. Acquisition parameters, drift, photostability, footprint and adhesion statistics of the 2-s and 5-s series, related to Figure 2 (CSV)
- Table S3. Acquisition parameters and photostability of the long-duration pulsed series, related to Figure 3 (CSV)
- Table S4. Continuous-wave power series, related to Figure 4 (CSV)
- Video S1. Raw pulsed-TIRF frames and cumulative single-molecule reconstruction of GFP-talin, related to Figure 2 (MP4)
- Video S2. One hour of pulsed TIRF of GFP-talin at one frame per second, related to Figure 3 (MP4)
- Data S1. Per-frame source data for Figures 2–4 and S1–S5, with the file manifest of the long-duration dataset (ZIP)

## Data and Code Availability

This study did not generate new unique reagents. Owing to a change of position, the authors no longer have direct access to the pulsed supercontinuum TIRF instrument described here; its components and configuration are documented in Methods, Figure 1 and Figure S3.

- Per-frame source sequired to reanalyse the data reported in this paper is available from the corresponding author upon request.

## Author Information

### Corresponding Author

Xian Hu – Department of Biosciences, University of Oslo, Oslo, Norway; Department of Microbiology (KLM-MIK), Division of Laboratory Medicine, Institute of Clinical Medicine, Faculty of Medicine, University of Oslo, Oslo, Norway; Mechanobiology Institute, National University of Singapore, Singapore.

### Author Contributions

Conceptualisation, F.M., M.P.S. and X.H.; methodology and instrumentation, F.M.; investigation, F.M. and X.H.; software, formal analysis and visualisation, F.M. and X.H.; resources and supervision, M.P.S. and O.B.; writing – original draft, X.H.; writing – review and editing, F.M., O.B. and X.H.

### Notes

X.H., F.M. and O.B. are inventors on patent application WO2021160723A1^30^ (microscope apparatus for extended-focus imaging), which has since lapsed; following the presentation of the technology at ELMI 2026, they are exploring its re-commercialisation, but no company has been formed and no funding has been secured. The remaining author, M.P.S., had no competing interests.

## Acknowledgments

We thank NorMIC IBV / Life Science Imaging Centre, University of Oslo, for providing the continuous-wave TIRF microscope used in this work, Dr. Jonathan Marcus Townson for discussion, Dr. Mingxi Yao for discussion and data curation, and Dr. Pan Meng for data curation. This work was supported by the Research Council of Norway (INFRASTRUKTUR 208310/F50, NorMIC Oslo; NALMIN-II infrastructure 322607), the Norwegian Cancer Society, and the National Research Foundation and Ministry of Education of Singapore (RCE R-714-105-004-135).

This work began as a side project of a talin mechanobiology collaboration with Michael P. Sheetz at the Mechanobiology Institute, National University of Singapore, where the pulsed-TIRF imaging reported here was performed in 2017 and 2018; the continuous-wave recordings were made in 2026 at NorMIC Oslo, University of Oslo. Edna (Xian Hu) and Felix worked with Mike for nearly 10 years while he was the director of the Mechanobiology Institute and received a tremendous amount of guidance and support from him. Mike died on 30 January 2025, and this manuscript is part of an effort to complete the projects he started with us.

## Declaration of Generative AI and AI-Assisted Technologies in the Writing Process

During the preparation of this work the authors used AI assistants (Claude and ChatGPT) to write and run image-analysis code, to prepare figures and to draft and edit the text. After using these tools, the authors reviewed and edited the content as needed and take full responsibility for the content of the publication.

## Supporting Information

**Figure S1.**
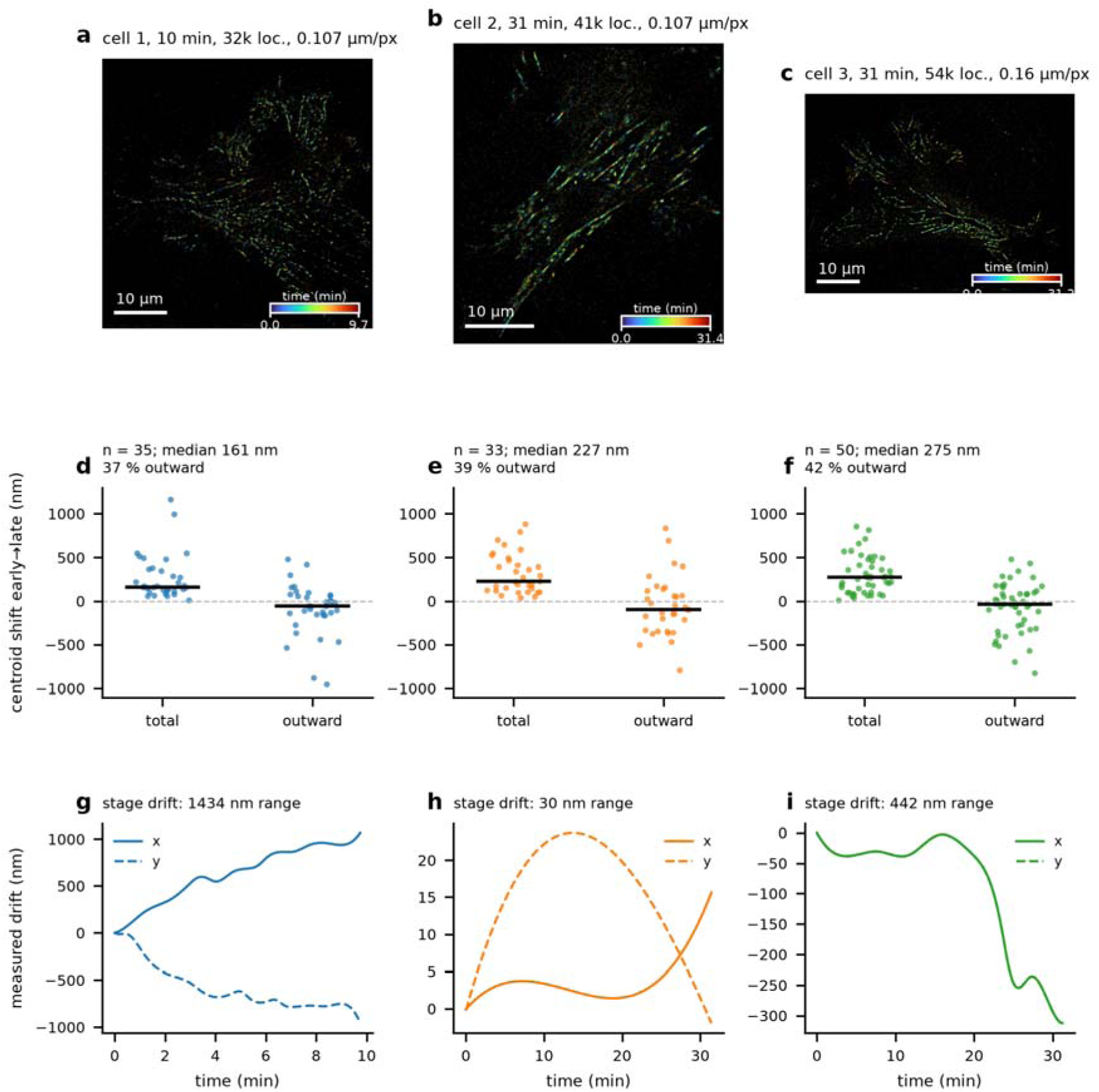
Pulsed-TIRF live-cell SMLM series: localisation maps, adhesion displacement and stage drift, related to Figure 2. (a–c) All localisations of cells 1–3 rendered at 20 nm per pixel and coloured by time of localisation (cell 1: 32,271 localisations, 9.8 min; cell 2: 41,992, 31.4 min; cell 3: 54,461, 31.2 min). Cells 1 and 2 were imaged at 107 nm per pixel, cell 3 at 160 nm per pixel. Scale bars 10 µm. (d–f) Per-adhesion centroid displacement between the first and last third of each series, total and outward component (n = 35, 33, 50 adhesions with ≥ 40 localisations in each third; bars, medians). (g–i) Measured stage drift (x solid, y dashed) for the three series; the correction applied to the data.

**Table S1.** Acquisition and analysis summary of the single-molecule series, related to Figure 2.

|  | **Cell 1** | **Cell 2** | **Cell 3** |
|---|---|---|---|
| Pixel size (nm) | 107 | 107 | 160 |
| Frames analysed / interval (s) / duration (min) | 283 / 2.07 / 9.8 | 874 of 901 / 2.10 / 31.4 | 901 / 2.08 / 31.2 |
| Cumulative 488 nm excitation (s) | 57 | 175 | 180 |
| Drift, total (nm) / residual r.m.s. (nm) | 1290 / 29 | < 25 / — | 313 / 15 |
| Localisations / per frame | 32,271 / 114 | 41,992 / 48 | 54,461 / 60 |
| Detection rate retained, last 20 vs first 20 frames | 79 % | 56 % | 104 % |
| Mean cell fluorescence retained | 78 % | 79 % | 90 % |
| PSF σ, median (nm) | 141 | 133 | 160 |
| Localisations inside adhesions / next-frame persistence inside, outside | 79 % / 20 %, 1 % | 79 % / 15 %, 1 % | 75 % / 17 %, 2 % |
| Adhesions in flow analysis / with significant slope | 36 / 12 | 35 / 20 | 52 / 36 |
| Significant adhesions: median \|v\| (nm min<sup>−1</sup>) / max / % retrograde | 58 / 197 / 67 | 18 / 47 / 60 | 19 / 70 / 61 |
| Centroid displacement early→late, median (nm) / interval (min) | 161 / 6.4 | 227 / 20.7 | 275 / 20.6 |
| Fraction of adhesions moving outward | 37 % | 39 % | 42 % |

Fluorescence retention is computed from the full-field Otsu cell mask for all three series; with the field cropped to the cell, the value for cell 3 is 107 %.

**Figure S2.**
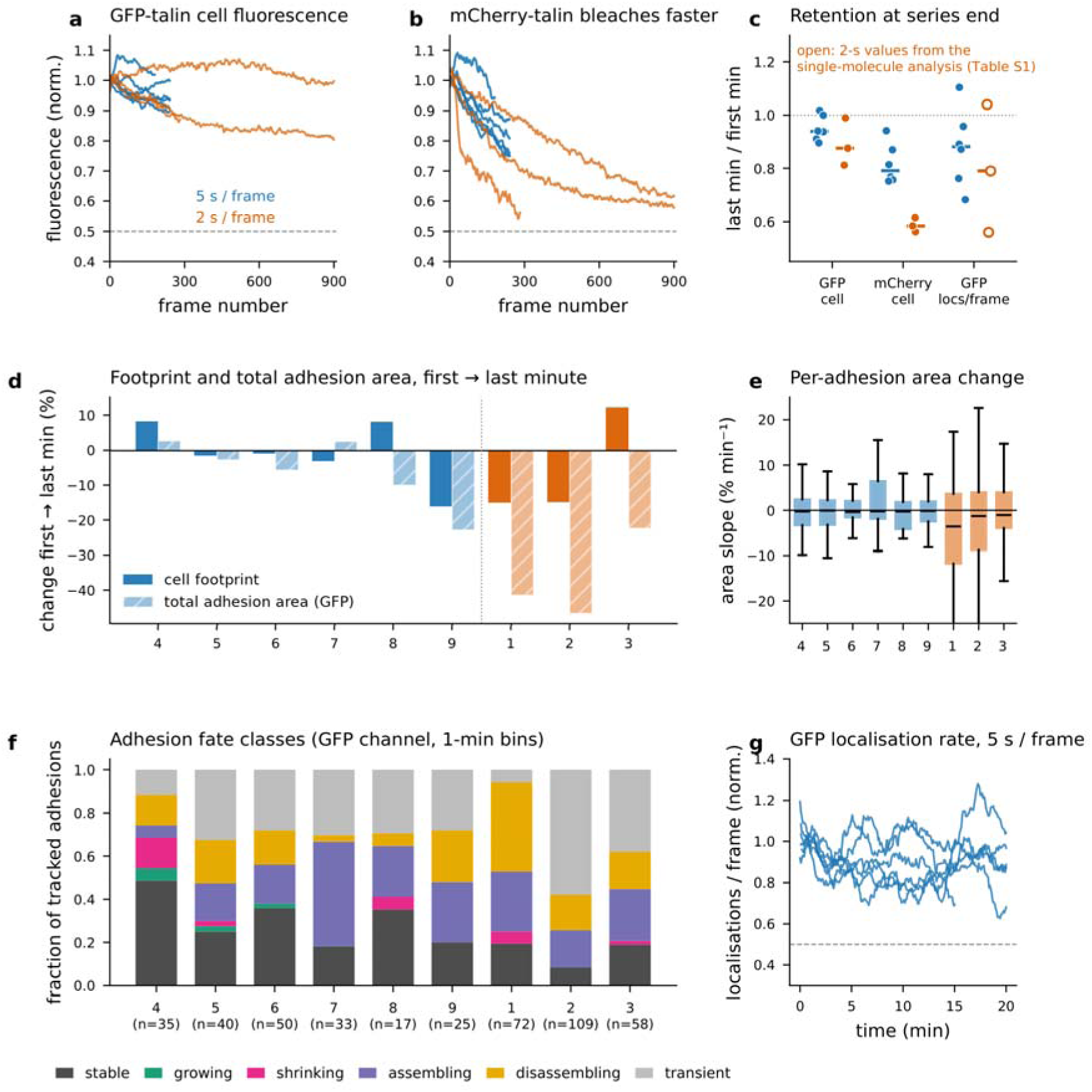
Cell health, adhesion turnover and photostability during 15–20 min of pulsed TIRF at 5-s intervals, compared with the 2-s series, related to Figure 2. (a, b) Background-subtracted mean cell fluorescence of GFP-talin (a) and mCherry-talin (b) versus frame number, normalised to the first minute (7-frame running mean); blue, 5-s series (cells 4–9); orange, 2-s series (cells 1–3, the recordings of Figure 2 and Figure S1). (c) Retention (last minute / first minute) of GFP and mCherry cell fluorescence and of the GFP localisation rate per cell; bars, medians. Filled symbols, values computed in this analysis; open symbols, the 2-s localisation-rate values taken from the single-molecule analysis (first 20 vs last 20 frames: 0.79, 0.56, 1.04; Table S1), as localisation was not repeated for the 2-s series here. (d) Change of cell footprint and of total GFP-talin adhesion area (bleach-normalised) between the first and last minute of each recording. (e) Distribution of per-adhesion area slopes (GFP channel, tracks ≥ 4 min; boxes, interquartile range; whiskers, 1.5 × IQR). (f) Fate classes of tracked adhesions (GFP channel): stable, growing, shrinking, assembling, disassembling and transient (see Methods); n, tracks per cell. (g) GFP localisations per frame (1-min running mean, normalised to the first 20 frames) for the six 5-s series.

**Figure S3.**
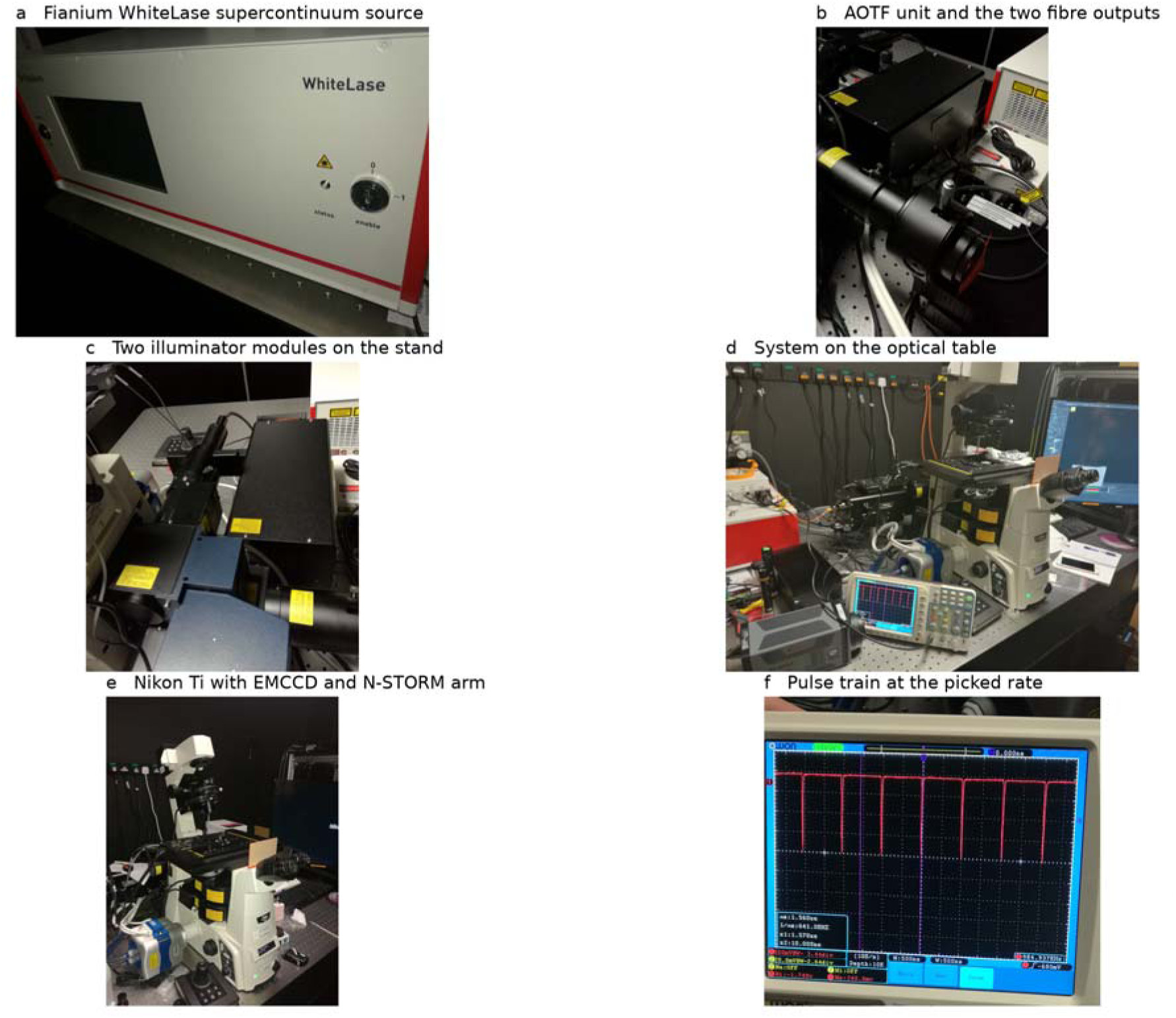
The instrument as built, related to Figure 1. Photographs of the system at the Mechanobiology Institute, National University of Singapore, taken during assembly (2016-07-28) and operation (2017-04-05 and 2017-05-20, the latter the day after the single-molecule series of Figure 2 were recorded); dates and times are from the camera metadata of each photograph. (a) The Fianium WL-SC450-8 WhiteLase picosecond supercontinuum source (2017-05-20). (b) The acousto-optic tunable filter and the two fibre outputs, mounted on the optical table ahead of the stand; the broadband input arrives from the source by the delivery fibre and leaves as the two polarisation-separated, spectrally windowed beams that feed the two illuminators (2016-07-28; corresponds to Figure 1a). (c) The two TIRF illuminator modules on the two layers of the Ti-E filter turret, with their free-space input beams entering from opposite sides of the stand at azimuths 90° apart (2016-07-28, five seconds after b; corresponds to Figure 1b). (d) The assembled system on the optical table during a recording session, with the oscilloscope monitoring the pulse train at the left of the stand and the acquisition computer at the right (2017-04-05). (e) The Nikon Ti-E stand with the EMCCD and the N-STORM illumination arm (2017-05-20). (f) The pulse train at the picked repetition rate, measured with a fast photodiode: negative-going pulses at 500 ns per division and 1 GS s⁻¹, with the instrument’s frequency counter reading 984.937 kHz (lower right), confirming the nominal 1 MHz used for every series reported here. The cursor read-out at lower left (Δx 1.560 µs, 1/Δx 641.0 kHz) reflects where the cursors happened to be placed and is not a measurement of the pulse rate. Photographed 2017-04-05, three minutes before d. The pulse duration (20–80 ps) is below the bandwidth of this oscilloscope and is taken from the manufacturer’s specification, not from this trace.

**Figure S4.**
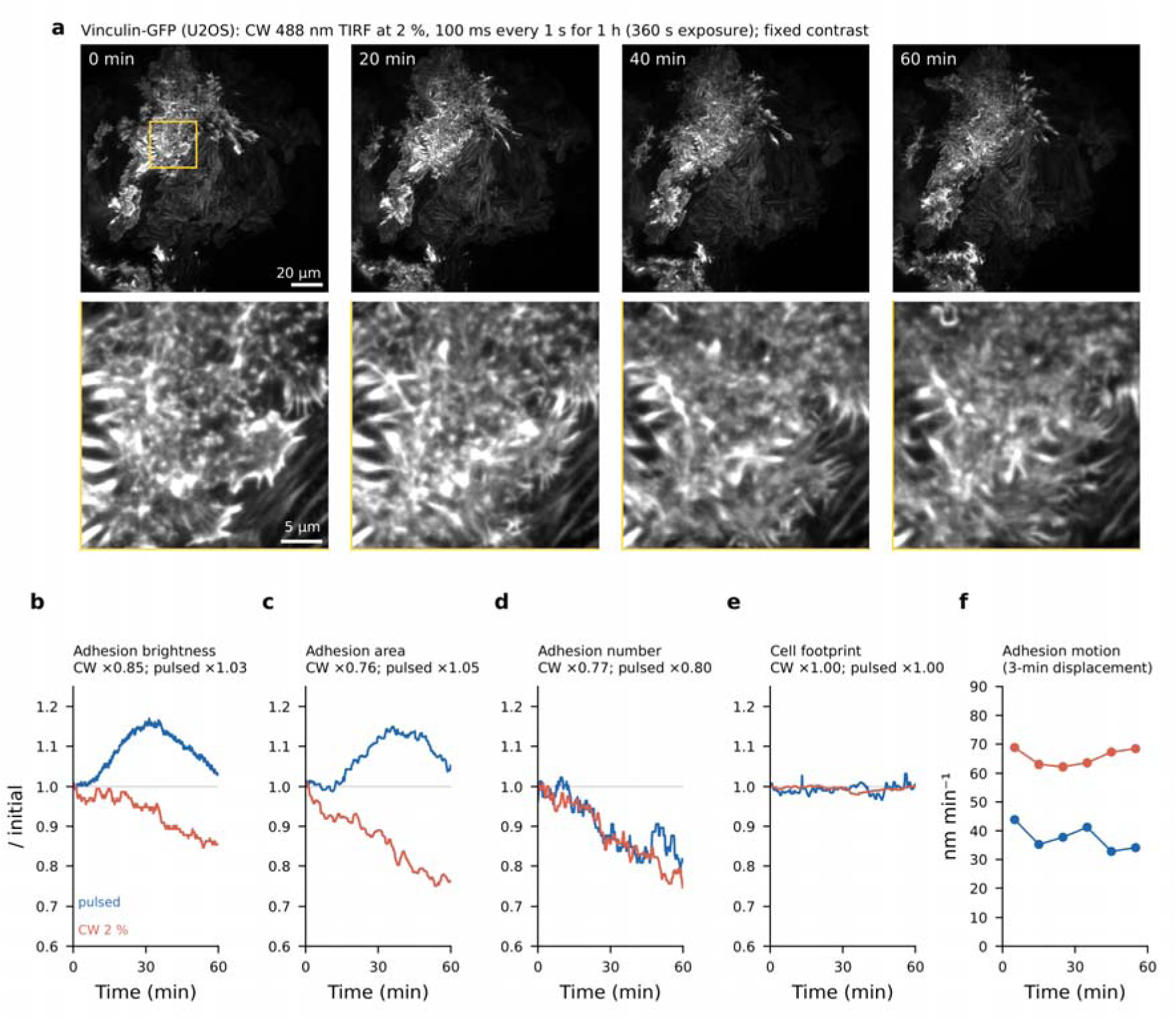
Under continuous-wave TIRF at a low laser setting the cell stays healthy for an hour but its adhesions dim; under pulsed TIRF at the same schedule they do not, related to Figure 4. (a) U2OS cell expressing vinculin-GFP imaged on a Nikon Ti2-E H-TIRF microscope (Apo TIRF 100×/1.49, Prime 95B sCMOS, 110 nm per pixel; H-TIRF angle 68.7°) with continuous-wave 488 nm excitation at a 2 % laser setting, 100 ms exposures every 1 s for 3600 frames (60 min, 360 s of cumulative excitation), shown after 0, 20, 40 and 60 min; bottom row, the boxed region enlarged. Each image is the mean of five consecutive frames, smoothed with a Gaussian of σ = 1 pixel, all at the contrast of the first image so that dimming is shown as recorded. Scale bars 20 µm (top) and 5 µm (bottom). (b–f) The read-outs of Figure 3 for this recording (red) and for the pulsed recording of Figure 3 (blue; MDA-MB-231, GFP-talin, the same schedule of 100 ms every 1 s for 60 min). (b) Adhesion brightness (99th percentile of the frame minus external background), normalised to the first 1 % of frames; running median of 15 frames. (c) Total area and (d) number of adhesions segmented with one fixed threshold for the whole hour, normalised to the first 2 min; running median of 7 time points. (e) Cell footprint, normalised to the first 2 min. (f) Median 3-min displacement rate of tracked adhesions in successive 10-min intervals. Panel titles give the end-to-start ratio of each read-out (Table 1). Analysis parameters were scaled from 160 to 110 nm pixels, and the adhesion threshold was set at the same fraction (0.158) of the initial adhesion brightness as for the pulsed recording (Methods). The two recordings differ in cell line, labelled protein, camera and instrument, and the laser power at the sample was measured for neither. The absolute rates in (f) are not comparable between the two recordings because adhesion density, and hence centroid noise, differs; what is compared is their constancy over the hour. Source data: Data S1.

**Figure S5.**
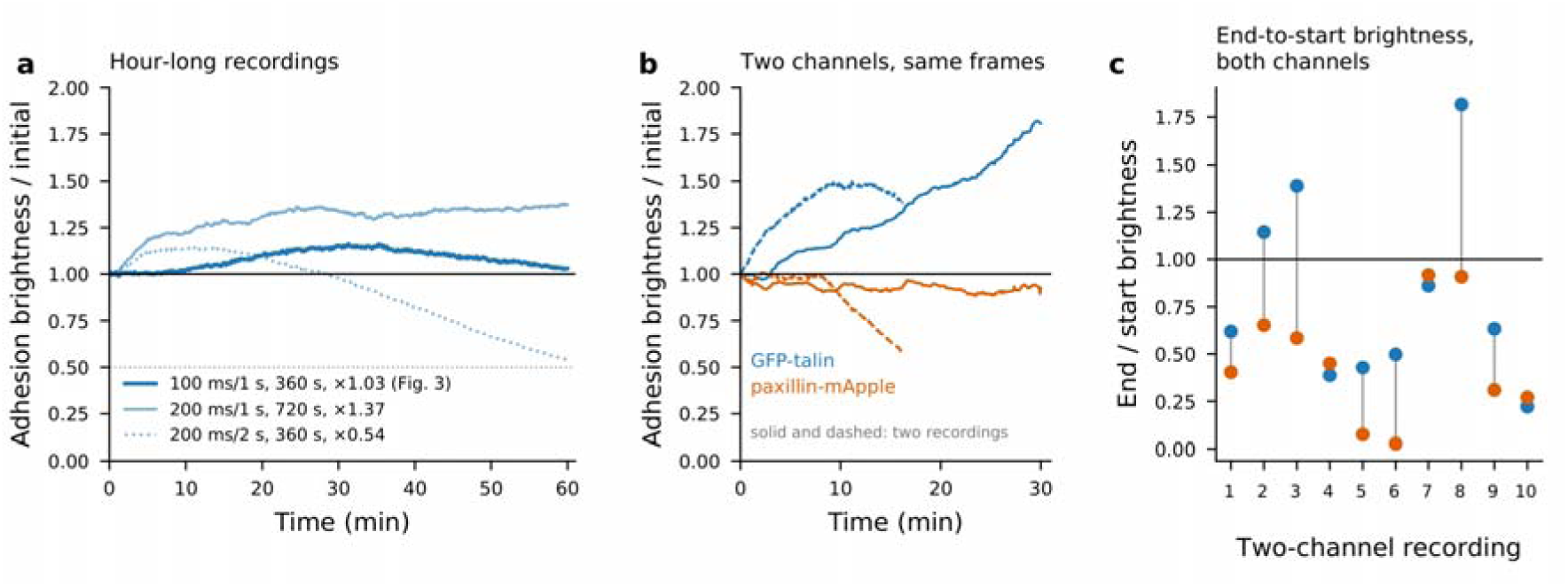
Hour-long and dual-colour series of the long-duration pulsed dataset, related to Figure 3. MDA-MB-231 cells co-expressing GFP-talin and paxillin-mApple, pulsed TIRF, 160 nm per pixel. (a) Adhesion brightness (99th percentile of the frame minus external background) normalised to the first 1 % of frames, for the recordings that lasted a full hour; the legend gives exposure and interval, cumulative excitation and the end-to-start ratio of each (AOTF settings in Methods). Solid line, unity; dotted line, 50 %. Running median of 9 frames. (b) The same quantity for the GFP-talin (blue) and paxillin-mApple (orange) channels of two dual-colour recordings, showing the GFP channel rising while mApple falls in the same frames. (c) End-to-start adhesion brightness of both channels in the dual-colour acquisitions, one vertical pair per recording; line at unity. Declining traces bound photobleaching from above and are not evidence of it (see Results); per-recording values in Table S3 and per-frame traces in Data S1.

### Supporting Notes

The notes give the full description of results that are summarised in the main text.

#### Note S1. Single-molecule series: acquisition, detection statistics, nature of the detections and drift

Cells expressing GFP-tagged talin (GFP at the N-terminal head; the construct also carries a C-terminal mCherry that was recorded in a second channel; Methods) were imaged on a Nikon Ti N-STORM TIRF microscope (100×/1.49 objective, 1.5× intermediate magnification, 107 nm pixels) with 20-nm bands centred at 488 nm and 561 nm selected by a pair of acousto-optic tunable filters from a picosecond supercontinuum laser (Fianium WL-SC450-8, 8 W total average output) at maximum transmission, acquired sequentially at each time point (200 ms GFP, 300 ms mCherry) on an EMCCD (EM gain 400) with a time-point interval of 2.07 s (Methods). Three series from the same day were analysed (Table S1): cell 1, 283 frames (9.8 min, 57 s cumulative excitation, 107 nm pixels); cell 2, 901 frames (31.4 min, 180 s, 107 nm pixels; 27 frames with transient intensity spikes or dips were discarded); cell 3, 901 frames (31.2 min, 180 s, 160 nm pixels, no intermediate magnification). All analyses use the GFP channel; unless stated otherwise, numbers in this section refer to cell 1, and the other cells are given in Table S1 and Figure 2e,f. Individual frames contained isolated, diffraction-limited spots on a diffuse background (Figure 2a; Video S1); a difference-of-Gaussians detector with a 4σ robust threshold followed by least-squares Gaussian fitting returned 32,271 localisations, 114 per frame, with a median fitted PSF width σ of 141 nm and a median spot integral of 3,440 camera units at EM gain 400 (Methods). The number of localisations per frame rose during the first minute, as the cell settled after the start of illumination, and then declined slowly; the mean of the last 20 frames was 79 % of the mean of the first 20 frames (Figure 2f). The mean GFP fluorescence of the cell followed the same course and retained 78 % of its initial value at the end of the series. In the two 31-minute series the number of detections per frame ended at 56 % (cell 2; 41,992 localisations, 48 per frame) and 104 % (cell 3; 54,461 localisations, 60 per frame) of its initial value, and the mean fluorescence at 79 % and 90 % (Figure 2f). Because talin turnover and adhesion remodelling also change both quantities — cell 3 shows no decline at all over 180 s of cumulative excitation — the observed losses are upper bounds on photobleaching. The second channel behaved similarly: the mCherry channel of cell 1 gave 10,727 localisations (38 per frame) with 90 % of the initial number of detections per frame retained, while its ensemble fluorescence fell to 62 %; across the series of the dataset the ensemble bleaching rate per second of exposure of mCherry was 1.2 times that of GFP (median; not significant), so mCherry bleached at least as fast as GFP, and in the 5-s series (Note S3) and the long-duration series (Note S4) the red fluorophores lost fluorescence faster than GFP. 79 % of all localisations fell inside adhesions segmented from the time-integrated density map, although these occupied only 34 % of the cell footprint.

Three observations indicate what a detection represents. First, detections are short. In cell 1 only 20 % of localisations inside adhesions, and 1 % outside (the chance level, 1 %), had a successor within 150 nm in the following frame, and across all series 87–96 % of linked events were confined to a single frame; the longest tracks that behaved as single emitters lasted 30–60 s. To resolve durations below the 2-s frame interval, one cell was streamed in the GFP channel alone at 0.33 s per frame for 10.0 min (1818 frames, frame-level duty cycle 100 %, 600 s of cumulative excitation). This series gave 102,818 detection events (57 per frame), 96 % of them confined to a single frame, corresponding to a mean bright-state duration of ≈ 0.35 s under a constant-hazard assumption (Methods). The number of detections per frame remained above 80 % of its peak for the full ten minutes; the fluorescence of the diffuse pool was 97 % of its initial value after 600 s of continuous exposure, while the brightest adhesion pixels declined with a half-time of about 10 min. Second, the emitting molecules are a small subset of the bound ones. The summed integrals of the fitted spots accounted for about 9 % of the background-subtracted adhesion fluorescence, and the adhesion fluorescence at any instant corresponded to about one emitting molecule per diffraction-limited area. Because both tags of the construct sit on the same molecule, the two channels provide an independent test: after channel registration (offset 50 nm in x, 62 nm in y), only 1.0 % of mCherry localisations coincided within 100 nm with a GFP localisation at the same time point, against 0.5 % for frame-shuffled controls. Third, the same sites emit repeatedly: pairs of detections within 40 nm recurred at time lags up to the 30 min of the recordings, compared with a control in which one copy of the coordinates was displaced by 250 nm (Methods). Together these observations indicate that a detection reports a brief bright period of an immobilised molecule rather than its presence, so that localisation counts are detection events and not molecule numbers. Across the series the loss of adhesion fluorescence was not ordered by cumulative excitation, as expected if it reflects adhesion turnover and remodelling more than photobleaching.

Cell 1, recorded shortly after the microscope was set up, drifted by 12.0 pixels (1.29 µm) over ten minutes, most of it in the first four minutes; cells 2 and 3, recorded later, drifted by < 25 nm and 313 nm over 31 minutes (Figure S1g–i). The drift was measured by phase cross-correlation of ten-frame averages with 20-fold sub-pixel upsampling, smoothed with a cubic spline and applied to every frame (Methods; Figure S1). Residual drift, measured independently by cross-correlating localisation histograms of 25-frame segments, was 0.27 pixels r.m.s. (29 nm) in cell 1 and 15 nm r.m.s. in cell 3, i.e. below the single-frame localisation precision.

#### Note S2. Stationary and mobile adhesions in the localisation data

Rendering all localisations colour-coded by acquisition time produced a map of the adhesion field in which each adhesion carries its own 10- or 31-minute history (Figure 2a–d; Figure S1a– c). Most adhesions appeared as mixed-colour structures, indicating that talin was localised at the same positions throughout the recording; a minority showed ordered colour gradients along their long axis (Figure 2b–d). To quantify this, adhesions were segmented from the time-integrated localisation density, and for each adhesion with at least 150 localisations spanning at least six minutes (n = 36) the localisations were projected onto the adhesion’s principal axis, oriented toward the cell centre, and position was regressed against time (Methods). The median along-axis speed over all adhesions was 8 nm min⁻¹ toward the cell centre (interquartile range −5 to 26 nm min⁻¹); 64 % of adhesions had a retrograde sign (Wilcoxon signed-rank p = 0.056). Twelve adhesions (33 %) had an along-axis speed significantly different from zero (p < 0.01). Of these, eight moved retrogradely, with speeds of 0.06–0.2 µm min⁻¹ (median 58 nm min⁻¹, maximum 197 nm min⁻¹) (Figure 2e; Table S1). In cells 2 and 3 the longer records resolved significant motion in a larger fraction of adhesions (20 of 35 and 36 of 52), but the speeds were lower: median |v| 18 and 19 nm min⁻¹, maxima 47 and 70 nm min⁻¹, again with ∼60 % retrograde and no significant directional bias (Figure 2e; Table S1). The mobile fraction observed in cell 1 was therefore not present in the two other cells, which were in a quieter state throughout their 31-minute recordings. An independent estimate from the displacement of adhesion centroids between the first and last third of each series gave median displacements of 161 nm over 6.4 min in cell 1 (25 nm min⁻¹), 227 nm over 20.7 min in cell 2 and 275 nm over 20.6 min in cell 3 (11 and 13 nm min⁻¹), with a slight inward (retrograde) median component and no net outward motion in any cell (37–42 % of adhesions moved away from the cell centre; p ≥ 0.10; Figure S1d–f). The speeds of the mobile fraction fall within the range reported for talin and vinculin flow in focal adhesions (0.1–0.3 µm min⁻¹; Hu et al., 2007; Case and Waterman, 2015) and match the retrograde speed of lamellar F-actin (∼2 nm s⁻¹ = 0.12 µm min⁻¹; Gardel et al., 2008), while being an order of magnitude slower than lamellipodial actin flow (∼25 nm s⁻¹). A plausible reading is that the mobile adhesions of cell 1 are those in which talin is coupled, through a slipping clutch, to the lamellar actin network, whereas the stationary majority are anchored to the substrate; establishing this, and the cell-to-cell variation in the mobile fraction, is a biological question beyond the scope of this methods report, which here only demonstrates that the measurement is possible at the required speed and duration. Two limitations apply to this interpretation. First, the measurement is population-based: the along-axis drift of the localisation cloud cannot distinguish the flow of talin molecules through a stationary adhesion from treadmilling or sliding of the adhesion itself. Second, only 16 % of localisations had a successor within 150 nm in the following frame (1 % expected by chance), so individual molecules were not tracked; at a 2 s interval a 0.1 µm min⁻¹ flow corresponds to 3 nm per frame, below the single-frame precision. Resolving the flow of molecules would require tracking individual emitters at shorter frame intervals, or speckle-type analysis; in the 0.33-s stream the bright periods (≈ 0.35 s) were too short to follow a molecule over the distances involved.

#### Note S3. Cell behaviour during 15–20 min of pulsed TIRF at 5-s intervals

To test whether the extended recordings perturb the cells, six further cells expressing GFP-talin-mCherry were imaged at 100 % laser power on both lines (488 nm, 330 ms; 561 nm, 300 ms, recorded in alternation) with one frame pair every 5.0 s for 15 min (181 frames; cell 4) or 20 min (241 frames; cells 5–9) at 160 nm per pixel (Figure S2; Table S2; cells are numbered as in Figure S2, cells 1–3 being the 2-s series). The 5-s interval is too sparse for localisation-based motion analysis but is well suited to a diffraction-limited read-out of cell behaviour, and the mCherry channel provides an independent, diffraction-limited image of the same adhesions. Stage drift was 21–157 nm over the recordings and was corrected as above; 2–7 frames per series showed brief illumination spikes and were excluded from intensity measurements.

Cell footprint, measured on 1-min averages of the GFP channel at a fixed threshold, changed by +8, −2, −1, −3, +8 and −16 % between the first and last minute of the six recordings (Figure S2d); five of the six cells exchanged only 2–10 % of their area between protrusion and retraction, and none rounded, blebbed or detached. The total area of GFP-talin adhesions (top-hat segmentation of 1-min averages, normalised for photobleaching by the cell-mean retention) changed by +3, −3, −6, +3, −10 and −23 % in the same cells, and the number of segmented adhesions was constant (Table S2). Tracking individual adhesions through the 1-min bins (n = 200 tracks of ≥ 4 min in the six cells) showed that the median per-adhesion area slope was between −0.37 and −0.06 % min⁻¹, that 18–49 % of tracked adhesions were stable (present throughout, no significant area change) and that only 4 of 200 adhesions grew significantly, whereas 2–16 new adhesions per cell assembled during the recording (Figure S2e, f). Cells 4 and 7 protruded and assembled new adhesions at their leading edges, i.e. the behaviour expected of unperturbed migrating fibroblasts; cell 9 retracted a 112 µm² rim and lost 23 % of its adhesion area.

Photobleaching was modest in the GFP channel: the background-subtracted mean cell fluorescence retained 0.90–1.02 (median 0.94) of its initial value after 181–241 frames, with the cytoplasmic pool (0.92–1.04) retaining more than the adhesion-integrated signal (0.72–0.89), as expected if bleached molecules in the exchanging cytosolic pool are replaced from the unilluminated volume whereas immobilised molecules in adhesions are not. mCherry bleached faster than GFP in every cell (cell mean 0.75–0.94, median 0.79; adhesions 0.59–0.80) (Figure S2a–c). The number of single-molecule detections per frame in the GFP channel, 30–82, was retained at 0.68–1.10 (median 0.88) of its initial value at the end of the recordings (Figure S2g), confirming that the persistence reported for the 2-s series does not depend on the frame interval.

For comparison, the three 2-s series (cells 1–3) were re-analysed with the identical morphology pipeline. Two of the three cells retracted by 15 % of their footprint, and all three lost 22–47 % of their GFP-talin adhesion area, with per-adhesion area slopes between −3.6 and −1.1 % min⁻¹ and only 8–19 % stable adhesions (Figure S2d–f); mCherry retention was 0.56–0.61 and GFP retention 0.81–0.99. The 2-s and 5-s recordings differ in exposure (200 vs 330 ms), EM gain and duty cycle (10 % vs 6.6 %) as well as in cell state, so with three and six cells the comparison does not identify the cause of the difference; it does indicate that at a 5-s interval the cumulative 488-nm excitation of 60–80 s per cell (equal to or greater than that of the 10-min series) is compatible with normal adhesion turnover, and the loss of adhesion area at 2-s intervals is a candidate phototoxic effect that these data can neither confirm nor exclude.

#### Note S4. The long-duration dataset

The series above are limited to 31 min and were optimised for single-molecule detection. To ask how far the photostability extends, we analysed an independent dataset recorded on the same microscope fourteen months later (2018-08-03/05/06) in MDA-MB-231 cells co-expressing GFP-talin and paxillin-mApple as two separate proteins: nineteen acquisitions of 132 to 5761 frames, with a 100 ms exposure (200 ms in two series) repeated at a fixed period of 1, 2, 5 or 10 s, i.e. an illumination duty cycle of 1–20 %, covering 4.4 min to 8.0 h (Table S3). At one frame per second this interval is too dense to be read as single molecules and the adhesions are continuously labelled, so these series were analysed as diffraction-limited images; the read-out is the brightness of the adhesions themselves, taken as the 99th percentile of each frame minus a background measured outside the cell, which is insensitive to changes in cell shape that confound a whole-field or fixed-mask mean (Methods).

Three recordings lasted a full hour, and we describe one of them in detail (Figure 3; Video S2). It was imaged with 100 ms exposures every 1 s for 3601 frames (360 s of cumulative excitation; AOTF line 1 at 50 %), with no drift correction needed (drift < 1 pixel over the hour). Its adhesion brightness ended at 1.03 of its starting value, after rising to about 1.15 at 30 min (Figure 3b). The adhesions remained individually resolved from the first frame to the last and remodelled throughout (Figure 3a): segmented on 5-frame means every 10 s, their total area ended at 1.05 of its starting value (150 µm² at the start) while their number fell to 0.80 (103 at the start), so that the mean adhesion grew from 1.5 to 1.9 µm² as adhesions elongated and merged (Figure 3c). The cell footprint within the field, measured on the diffuse GFP-talin signal above the per-frame background, was unchanged (1.00; 0.98–1.02 over a two-fold range of threshold; Figure 3d); the cell extends beyond the right and lower edges of the field, so this is the area inside the field rather than the whole cell. Tracking the centroids of the segmented adhesions gave 196 tracks of at least 3 min (median 10 min); 37 % of them moved faster than the 95th percentile (32 nm min⁻¹) of the same tracks with their time points shuffled, at 32–223 nm min⁻¹ (median 57 nm min⁻¹), and 72 % of these mobile adhesions moved toward the cell centre (Figure 3e, f). These speeds are comparable to those resolved by single-molecule localisation in cell 1 of the 2017 series (retrograde adhesions at 0.06–0.2 µm min⁻¹, median 58 nm min⁻¹; Figure 2e), in a different cell line and with a different read-out, and the rate of adhesion displacement did not slow over the hour (median 3-min displacement rate 33–44 nm min⁻¹ in each 10-min interval; Figure 3g), as it would if the cell were progressively photodamaged. They also lie within the range reported for focal-adhesion sliding in unperturbed cells, toward its slow end: 0.12 ± 0.08 µm min⁻¹, directed toward the cell centre, for focal adhesions of stationary fibroblasts (Smilenov et al., 1999), and a mean of 19 µm h⁻¹ (0.3 µm min⁻¹) for small focal contacts translocating centripetally (Zamir et al., 2000). Because published values differ several-fold with cell type, adhesion type, substrate and the criterion for a moving adhesion, and because we have no unilluminated control for this cell, the comparison shows that the adhesions move at a plausible speed and in the expected direction; the evidence that the illumination did not slow them is the constancy of the speed over the hour. For this cell, then, an hour of imaging at one frame per second left the GFP signal, the adhesion area, the cell footprint and the rate of adhesion movement where they started.

The other two hour-long recordings bracket it (Figure S5a). In the first, 3601 frames at 1 s intervals with a 200 ms exposure — 720 s, i.e. 12 min, of cumulative excitation, with AOTF line 1 at 100 % — the adhesion brightness at the end of the hour was 1.37 times its starting value; in the second (1801 frames at 2 s, 200 ms, 360 s cumulative; all four 488-nm AOTF lines at 100 %) it was 0.54. The longest recording in the set, 5761 frames over 8.0 h at a 2 % duty cycle (576 s cumulative), ended at 0.46. Across the whole dataset the end-to-start ratio was not ordered by cumulative excitation: the largest dose, 720 s, gave the highest retention of any hour-long recording, while recordings receiving 36–50 s span 0.03 to 1.82 (Table S3).

A declining trace cannot by itself be attributed to photobleaching, because cell stress, focus or TIRF-angle drift, adhesion disassembly, protein turnover and migration out of the evanescent field all reduce the signal; each trace is therefore an upper bound on bleaching, as for the series above. The reverse is not true: no mechanism restores a bleached fluorophore, so a cell whose adhesion brightness is unchanged after an hour of excitation establishes that photobleaching was negligible in that recording, and the cells that declined are uninformative rather than contradictory. The ten dual-colour acquisitions add an internal control, since both fluorophores in a given cell experience the same illumination schedule, focus and cell state: paxillin-mApple ended below its starting brightness in 10 of 10 cells (0.03–0.92), whereas GFP-talin did so in 7 of 10 and ended brighter in three (1.14, 1.39 and 1.82), and in two cells the GFP trace rose while the mApple trace in the same frames fell (Figure S5b, c). A change of focus or of the evanescent field would move both channels together, so the divergence localises the loss to fluorophore photophysics. This control is weaker than the dual-tagged construct used above, because talin and paxillin are different proteins with different turnover and adhesion residence; it constrains optical and cell-state artefacts but not protein-specific exchange. Taken with the 2-s and 5-s series, the dataset shows that GFP-talin tolerates at least an hour of continuous pulsed TIRF at a 10–20 % duty cycle without net loss of adhesion fluorescence, in a second cell type, whereas the co-expressed red fluorophore, mApple, did not share this persistence. As for the shorter series, attributing this to pulsing rather than to the fluorophore, to the low frame-level duty cycle or to exchange would require a continuous-wave recording on the same instrument and specimen, which does not exist; the continuous-wave recordings described next were made eight years later on a different microscope and specimen.

#### Note S5. The continuous-wave power series

Eight of the nine continuous-wave series used the 100 ms per 1 s schedule, for 3.4–60 min (20– 360 s of cumulative excitation); the ninth, at 1 %, was streamed at 0.117 s per frame for 2.2 min (112 s). The laser setting was constant within each recording. At 1–5 % the cells kept most of their GFP signal: footprint-integrated fluorescence ended at 0.94–1.01 of its starting value in all four series, and none of the four cells retracted (footprint area 0.96–1.03 at the end of the recording). At 8–10 % the outcome was different in all five cells (Figure 4a, d, e). Footprint fluorescence stayed above 0.8 for the first 16–57 s of exposure and then fell, reaching half its starting value after 25–50 s in four of the five cells; it ended at 0.02–0.20 in those four and at 0.74 in the fifth (62 s of exposure). After 45 s of exposure, the footprint fluorescence was 0.97–1.06 at 1–5 %, 0.58 and 0.90 at 8 % and 0.19–0.37 at 10 %, against 1.02–1.17 in the three hour-long pulsed series (Figure 3, Figure S5; Figure 4e). The cells retracted over the same period: the two isolated cells that could be segmented throughout (8 %) ended at 0.66 and 0.83 of their starting footprint area (Figure 4f), and in the hour-long 10 % series the spread cells had contracted into compact bodies by 20 min (Figure 4a). Because retraction lifts fluorescence out of the evanescent field, the loss of TIRF signal at 8–10 % combines photobleaching with the loss of cell–substrate contact, and these data do not partition it between the two.

The loss rose much more steeply than the laser setting. At 10 %, footprint fluorescence halved after 25–29 s of exposure; if the loss were proportional to setting × exposure, the 2 % recording, made at a similar TIRF angle (68.7° against 67.6°), would have halved after 125–145 s, whereas it retained 0.94 after 360 s. The 5 % series were made at a steeper TIRF angle (72.6°, nominal penetration depth 77 nm) and gave dim images (signal-to-noise ratio 9.5–9.8), so where between 2 and 8 % the transition lies is not determined. On this microscope, then, the useful duration of continuous-wave GFP imaging was limited by the laser setting rather than by cumulative exposure, and high settings shortened it by more than an order of magnitude through bleaching and cell retraction together. The laser power at the sample was not measured at any setting, and the TIRF angle varied between 65.3° and 72.6° between sessions, so these recordings describe continuous-wave GFP TIRF on a standard instrument and do not compare it with pulsed excitation at matched irradiance.

**Table S5.** Key resources: cell lines, constructs, reagents, instruments and software.

| **REAGENT or RESOURCE** | **SOURCE** | **IDENTIFIER** |
|---|---|---|
| **Experimental models: Cell lines** |  |  |
| Human: MDA-MB-231 | Laboratory of Michael P. Sheetz, Mechanobiology Institute, National University of Singapore | RRID:CVCL_0062 |
| Human: U2OS | Laboratory of Michael P. Sheetz, Mechanobiology Institute, National University of Singapore | RRID:CVCL_0042 |
| **Chemicals, peptides, and recombinant proteins** |  |  |
| jetOPTIMUS DNA transfection reagent | Polyplus (Sartorius) | Cat# 201000001 |
| **Recombinant DNA** |  |  |
| GFP-talin-mCherry (GFP at the N-terminal head, mCherry at the C-terminal rod) | Margadant et al., 2011; Hu et al., 2016 | N/A |
| GFP-talin | Laboratory of Michael P. Sheetz, Mechanobiology Institute, National University of Singapore | N/A |
| Paxillin-mApple | Laboratory of Michael P. Sheetz, Mechanobiology Institute, National University of Singapore | N/A |
| Vinculin-GFP | Laboratory of Michael P. Sheetz, Mechanobiology Institute, National University of Singapore | N/A |
| **Deposited data** |  |  |
| Raw image series and per-frame source data | This paper | BioImage Archive (to be deposited during peer review); Data S1 |
| **Software and algorithms** |  |  |
| Python 3.13 | Python Software Foundation | https://www.python.org |
| nd2 0.11.3 | Python Package Index | https://pypi.org/project/nd2 |
| tifffile 2026.6.1 | Python Package Index | https://pypi.org/project/tifffile |
| scikit-image 0.26 | Python Package Index | https://scikit-image.org |
| SciPy, NumPy, Matplotlib | Python Package Index | https://scipy.org; https://numpy.org; https://matplotlib.org |
| ND2-to-OME-TIFF ImageJ plugin | This paper | https://github.com/evildna/Nikon-ND2-to-OMETIFF-ImageJ-plugin |
| Drift-correction ImageJ plugin for SMLM time series | This paper | https://github.com/evildna/DriftCorrectionForSMLMImageJPlugin |
| Simple_SMLM_Localizer_2D_TIRF ImageJ plugin for single-molecule localisation | This paper | https://github.com/evildna/Simple_SMLM_Localizer_2D_TIRF |
| PulsedTIRF_analysis Python analysis code | This paper | Zip file with this submission; Zenodo DOI and https://github.com/evildna/PulsedTIRF_analysis at publication |
| **Other** |  |  |
| Picosecond supercontinuum laser WL-SC450-8 with pulse picker | Fianium | WhiteLase WL-SC450-8 |
| Inverted microscope Ti-E with N-STORM TIRF illuminator | Nikon | N/A |
| HP Apo TIRF 100×/1.49 objective | Nikon | N/A |
| Evolve Delta EMCCD camera | Photometrics | N/A |
| Inverted microscope Ti2-E with H-TIRF module and Apo TIRF 100×/1.49 objective | Nikon | N/A |
| Prime 95B sCMOS camera | Photometrics | N/A |
| Continuous-wave laser combiner LightHUB+ (LightHUBplus N) with 405/488 nm lasers, 60/60 mW (maximum output 60 mW at 488 nm) | Omicron-Laserage Laserprodukte, Rodgau, Germany | MOM50201 LH+APC |
| 35-mm glass-bottom dish, 14-mm microwell, No. 1.5 coverglass | MatTek | Cat# P35G-1.5-14-C |
| µ-Dish 35 mm, high, Glass Bottom (No. 1.5H coverslip) | ibidi | Cat# 81158 |

### Titles and legends of the Supporting Information files provided separately

**Table S2. Acquisition parameters, drift, photostability, footprint and adhesion statistics of the 2-s and 5-s series, related to Figure 2**. Provided as a separate file.

**Table S3. Acquisition parameters and photostability of the long-duration pulsed series, related to Figure 3**. Provided as a separate file: per recording and channel, frame count, interval, exposure, duty cycle, duration, cumulative excitation, background fraction, and the end-to-start ratio of both the adhesion brightness and the fixed-mask cell mean. Per-frame traces are in Data S1.

**Table S4. Continuous-wave power series, related to Figure 4**. Provided as a separate file: per recording, laser setting, pixel size, frame count, duration, cumulative exposure, single-frame adhesion signal-to-noise ratio, end-to-start ratios of adhesion brightness and footprint fluorescence, cumulative exposure at which footprint fluorescence fell below 0.8 and 0.5, and footprint area at the end and minimum.

**Video S1. Raw pulsed-TIRF frames and cumulative single-molecule reconstruction of GFP-talin, related to Figure 2**. Left, drift-corrected raw pulsed-TIRF frames of GFP-talin (fixed contrast); right, cumulative SMLM reconstruction coloured by time of localisation, with the localisations of the current frame circled. 283 frames, 2.07 s interval, played at 15 frames s⁻¹. Scale bars 10 µm.

**Video S2. One hour of pulsed TIRF of GFP-talin at one frame per second, related to** **Figure** 3. The recording of Figure 3 (3601 frames, 100 ms every 1 s, 360 s of cumulative excitation). Every fifth frame is shown, each the mean of five consecutive frames, smoothed with a Gaussian of σ = 0.7 pixel, at a fixed contrast set on the first frames; 720 frames played at 30 frames s⁻¹ (150× real time). Scale bar 10 µm.

**Data S1. Per-frame source data, related to Figures 2–4 and S1–S5.** Comma-separated tables behind each quantitative panel, the drift and per-adhesion tables of the single-molecule series, the per-frame traces and file manifest of the long-duration dataset and the acquisition metadata of the continuous-wave series; a README maps each file to its figure panel.

